# Structural Basis for Activation of HRI by the DELE1 C-terminal domain

**DOI:** 10.64898/2026.09.15.751698

**Authors:** Olawale G. Raimi, Rupam Bhattacharjee, Min Cao, Shannon Richardson, Andrew C. H. Liu, Vanesa Vinciauskaite, Carine De Marcos Lousa, Miratul M.K. Muqit, Glenn R. Masson

**Affiliations:** Division of Cancer Research, Faculty of Health, University of Dundee, Dundee, DD1 9SY, United Kingdom; MRC Protein Phosphorylation and Ubiquitylation Unit, Faculty of Life Sciences, University of Dundee, Dundee, DD1 5EH, United Kingdom; UK Dementia Research Institute Parkinson’s Research Centre at University of Edinburgh, Edinburgh, United Kingdom; Institute for Neuroscience and Cardiovascular Research, School of Neurological and Cardiovascular Sciences, College of Medicine and Veterinary Medicine, The University of Edinburgh, United Kingdom

## Abstract

Haem-Regulated Inhibitor (HRI, aka EIF2AK1) is one of four stress-sensing kinases that phosphorylate eIF2α as part of the integrated stress response (ISR). HRI was first characterized as a haem-sensing kinase where it is inhibited when bound to haem. Recent studies have revealed another role for HRI in sensing mitochondrial stress via the mitochondrial protein DELE1. Upon stress, DELE1 is cleaved, and its C-terminal fragment (DELE1^CTD^) is released from the mitochondria to the cytoplasm, where it interacts with HRI, to trigger the ISR. This pathway is critical for mitochondrial quality control and neuronal health. Here, we perform biophysical analysis to demonstrate that purified recombinant DELE1^CTD^ binds with high affinity to the N-terminal region of HRI, competing with and displacing haem to activate HRI and pointing to a shared binding site. Using Hydrogen Deuterium Exchange Mass Spectrometry (HDX-MS), we map the binding footprint of DELE1^CTD^ on HRI and confirm that it overlaps with the haem-binding site. These findings support a simple activation mechanism: DELE1^CTD^ activates HRI by excluding inhibitory haem from its binding site.

## Introduction

Haem-regulated inhibitor (HRI, also known as EIF2AK1) is a haem-sensing protein kinase that exists in an autoinhibited state when bound to haem (Farrell *et al*, 1977; Yang *et al*, 1992). Upon the loss of haem, HRI is activated and subsequently auto-phosphorylates and phosphorylates the eukaryotic initiation factor 2α (eIF2α), activating the integrated stress response (ISR) resulting in a global arrest of protein translation, with the exception of a subset of stress responsive elements (Chen, 2025; Singh *et al*, 2025; Kanta *et al*, 2025; Sekine *et al*, 2023; Fessler *et al*, 2020; Yang *et al*, 2023; Guo *et al*, 2020; Zebrucka *et al*, 2016). HRI was, until relatively recently, believed to be found only in erythroid progenitor cells (Crosby *et al*, 1994), but it has since been discovered that HRI is ubiquitously expressed with a highly conserved role in sensing mitochondrial damage (Abdel-Nour *et al*, 2019; Singh *et al*, 2025; Sekine *et al*, 2023). Mitochondrial dysfunction is a major hallmark of multiple diseases, including neurodegenerative disorders such as Parkinson’s disease, where our groups and others recently demonstrated that HRI activation from damaged mitochondria inhibits PTEN-induced kinase 1 (PINK1) – PARKIN ubiquitin-dependent mitophagy (Singh *et al*, 2025; Yang *et al*, 2026). The critical signalling factor for mitochondrial HRI activation is DAP3-binding cell death enhancer 1 (DELE1), a protein localised within the mitochondrial inner membrane, but which under mitochondrial stress (that can be induced by depolarisation or the inhibition of oxidative phosphorylation) is cleaved by the inner mitochondrial membrane metalloprotease OMA1, producing a C-terminal fragment that is released and accumulates in the cytosol (Yang *et al*, 2023; Guo *et al*, 2020; Sekine *et al*, 2023; Fessler *et al*, 2020; Bi *et al*, 2024; Bora *et al*, 2025; Yang *et al*, 2026).

The structure of the oligomeric C-terminal fragment of DELE1 (DELE1^CTD^) was recently solved by cryo-EM, detailing the orientation of 11 Tetratricopeptide repeat (TPR) helices (Yang *et al*, 2023), with the rest of the protein predicted to be largely unstructured. A highly truncated structure of the kinase domain of HRI has recently been solved (although missing the large, disordered kinase insert (**See Figure 1**)) in high resolution (Rajasekaran *et al*, 2026). This structure is missing the N-terminal haem binding domain of HRI, which, primarily through cell-based studies, has previously been implicated in DELE1-mediated activation of HRI (Fessler *et al*, 2020). Previously, our group has used biochemical assays and structural Hydrogen Deuterium Exchange Mass Spectrometry (HDX-MS) to glean insight into how HRI is directly regulated by haem, and these data revealed that large structural rearrangements across the entire protein accompany haem binding and HRI autoinhibition, although the haem binding domain appeared to be the primary site of interaction (Kanta *et al*, 2025).

**Figure 1:**
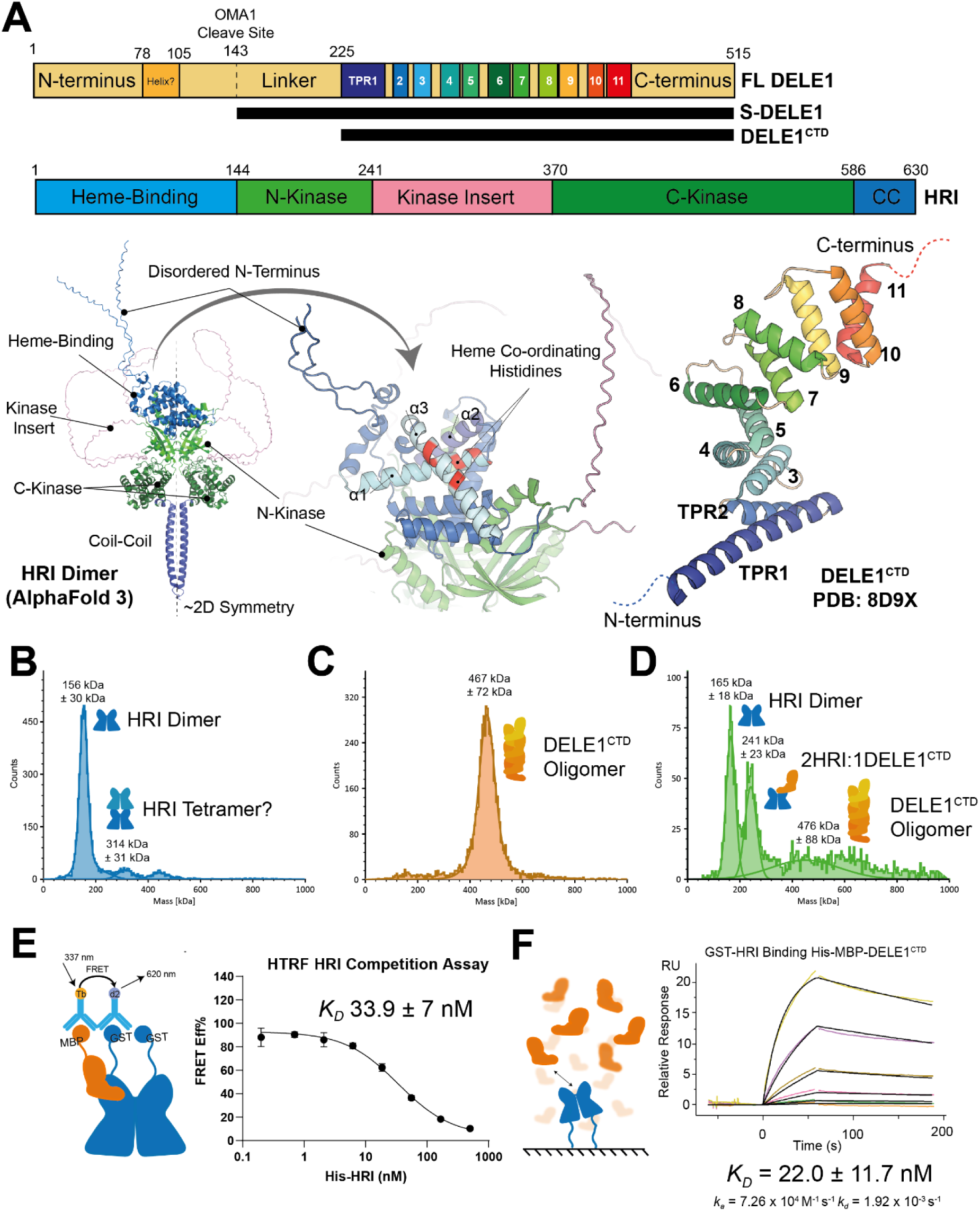
HRI binds directly to DELE1^CTD^. **(A)** Schematic of the constructs used in this study include full-length DELE1 (FL DELE1, residues 1-515), S-DELE1 (Residues 143-515), DELE1^CTD^ (225-515) and full-length HRI (residues 1-630). TPR = Tetratricopeptide repeat sequence. AlphaFold structure model of HRI and cryo-EM structure of DELE1^CTD^ (adapted from PDB: 8D9X). (**B**) Mass photometry of 100 nM His-HRI showing a dimeric (predicted 150 kDa) and possible 4-mer peak (predicted 300 kDa). (**C**) Mass photometry of 100 nM His-MBP- DELE1^CTD^ showing an oligomeric state (predicted monomer = 72 kDa) (**D**) Mass photometry of 100 nM His-MBP- DELE1^CTD^/His-HRI showing the formation of a novel peak at mass 241 kDa, indicating an HRI dimer bound to a single DELE1^CTD^ monomer. All mass photometry histograms are representative single data collections, while masses indicated are means and standard deviations from three independent measurements. (**E)** Schematic of the TR-FRET assay of GST-HRI and His-MBP-DELE1^CTD^. After stable complex formation, His-HRI is titrated into the complex to compete away the DELE1^CTD^, producing an HRI binding affinity measurement of 33.9 nM. Data n = 3, individual points are means with standard deviation, some error bars are smaller than the data points. **(F)** Representative SPR sensorgram. Immobilised HRI was bound to DELE1^CTD^ in the mobile phase. An affinity of binding of 22 nM was measured, in agreement with the *in-solution* measurement achieved in (E). Experiment was repeated n = 4, binding constants shown are the mean of biological repeats.

The mechanism of how DELE1 binding facilitates HRI activation remains unknown. It is unclear whether the interaction is direct, whether it requires accessory factors and whether it can occur in the presence of haem, and neither the stoichiometry nor the binding interface has been defined. In this study, using biophysical analysis of purified recombinant proteins, we show that DELE1^CTD^ directly binds to HRI with nanomolar affinity and can displace haem, which in turn leads to activation of HRI. We further show that a single monomer of DELE1^CTD^ directly interacts with dimeric HRI and using structural HDX-MS map the interaction of DELE1^CTD^ to the disordered N-terminus of HRI and α-helices that are responsible for haem binding. This interaction is sufficient to directly overcome haem-induced inhibition of HRI, suggesting a simple model of how DELE1^CTD^ promotes activation of HRI under mitochondrial stress.

## Results

### HRI binds directly to DELE1^CTD^ with high affinity

In previous work, the interaction of DELE1 with HRI has been suggested by largely cell-based studies (Singh *et al*, 2025; Fessler *et al*, 2020; Sekine *et al*, 2023). To investigate whether HRI and DELE1 directly interact, N-terminal His or GST tagged full-length human HRI (amino acid (aa) residues 1-630) His-MBP tagged full-length DELE1 (aa residues 1-515), His-MBP tagged DELE1 fragment corresponding to the cleaved species (aa residues 143-515 (S-DELE1)); and a His-MBP tagged C-terminal DELE1 fragment (residues 225-515 (DELE1^CTD^ (Yang *et al*, 2023)) were expressed in *E. coli* and purified (**See Figure 1A and Supplementary Figure 1)**. As previously reported, we observed that recombinant HRI purified as an autophosphorylated dimer (Kanta *et al*, 2025), and as others have reported, both recombinant DELE1^CTD^ and S-DELE1 innately underwent oligomerisation following expression *in vitro,* whilst full-length DELE1 purified predominantly as a monomer (Yang *et al*, 2023) (as determined by calibrated size exclusion chromatography) **(Supplementary Figure 1).**

Initially, we determined a direct interaction between HRI and DELE1^CTD^ using mass photometry **(Figure 1B-D)**. Mass photometry analysis of 100 nM HRI determined an mass of ∼156 ± 30 kDa (predicted mass of a His-HRI monomer = 75 kDa) **(Figure 1B)**, while 100 nM DELE1^CTD^ produced an oligomeric peak of ∼467 ± 72 kDa (predicted His-MBP DELE1^CTD^ mass 72 kDa **(Figure 1C)**) suggesting a DELE1^CTD^ oligomer averaging approximately 6 monomers of DELE1^CTD^. However, upon incubation of 100 nM HRI with equimolar DELE1^CTD^, we strikingly observed a third peak with an average mass of 241 ± 23 kDa, suggesting a complex of dimeric HRI bound to a single monomer of DELE1^CTD^ **(Figure 1D)** (2HRI: 1DELE1^CTD^ = 222 kDa).

To gain a quantitative assessment of the interaction, we next established a time-resolved Förster resonance energy transfer (TR-FRET) assay of HRI and DELE1^CTD^ binding (**Figure 1E)**. Following optimisation and reciprocal titration experiments **(Supplementary Figures 2A-F)**, addition of 5 nM His-MBP-DELE1^CTD^ or 3 nM GST-HRI with fluorescently Anti-MBP (donor) or Anti-GST (acceptor) labelled antibodies, produced a robust and stable TR-FRET signal, suggestive of GST-HRI:DELE1^CTD^ complex formation. Consistent with this, under similar conditions, we did not observe FRET signal following incubation of GST-ubiquitin with His-MBP-DELE1^CTD^ in the presence of donor and acceptor antibodies **(Supplementary Figure 2E/F).** We next performed competition experiments via titration of increasing amounts of His-HRI and observed concentration-dependent ablation of FRET signal consistent with disruption of GST-HRI:DELE1^CTD^ complex formation. This enabled determination of the binding affinity (*K_D_*) of HRI to DELE1^CTD^ at 33.9 ± 7 nM (**Figure 1E)**. We next investigated what impact HRI (auto)phosphorylation may have on HRI’s interaction with DELE1. Addition of (auto)phosphorylated HRI (produced as previously described (Kanta *et al*, 2025)) to DELE1^CTD^ did not impact FRET signal, suggesting that HRI (auto)phosphorylation does not affect HRI:DELE1^CTD^ complex formation (**Supplementary Figure 2G)**.

We next developed an orthogonal surface plasmon resonance (SPR) assay to evaluate direct HRI:DELE1^CTD^ binding and complex formation **(Figure 1F)**. Using SPR, we immobilised GST-HRI on the chip surface before flowing His-MBP-DELE1^CTD^ in the mobile phase. Using an 8 point 3-fold serial dilution curve of His-MBP-DELE1^CTD^ and a top concentration of 600 nM, we observed that DELE1^CTD^ bound to HRI with a *K_D_* of 22.0 ± 11.7 nM. Stoichiometry analysis using theoretical R_max_ values favoured a single DELE1^CTD^ protein binding to each HRI dimer, although the binding of two DELE1^CTD^ proteins to each HRI dimer could not be excluded. The reciprocal experiment *i.e.* immobilised DETE1 and mobile HRI, we were unable to detect full-length DELE1 interaction with HRI while still observing HRI interaction with DELE1^CTD^ (**Supplementary Figure 3).**

### DELE1^CTD^ and Haem are competitive and mutually exclusive binders of HRI

We reconstituted HRI catalytic kinase activity by using ADP-GLO and incubating 50 nM recombinant human HRI with 5 µM of recombinant human eIF2α (EIF2S1) and 250 µM ATP, and consistent with our previous study (Kanta *et al*, 2025), we observed that the haem analogue, hemin, inhibited HRI kinase activity with an IC_50_ of 3.1 ± 2.1 μM - although it should be noted we did not observe complete inhibition, in line with many other reports **(Figure 2A)** (Ricketts *et al*, 2022). We next determined the binding affinity (*K_D_*) of hemin for HRI, using SPR with an immobilised GST-HRI with a 10-point dilution series of hemin as an analyte **(Figure 2B)**. From this data we were able to measure a *K_D_* of 5.1 ± 1.1 µM with a 1:1 stoichiometry (*i.e.* a single hemin molecule bound to single polypeptide of HRI) (as determined using a theoretical Rmax), in good agreement with the kinase activity assay **(Figure 2A)**.

**Figure 2:**
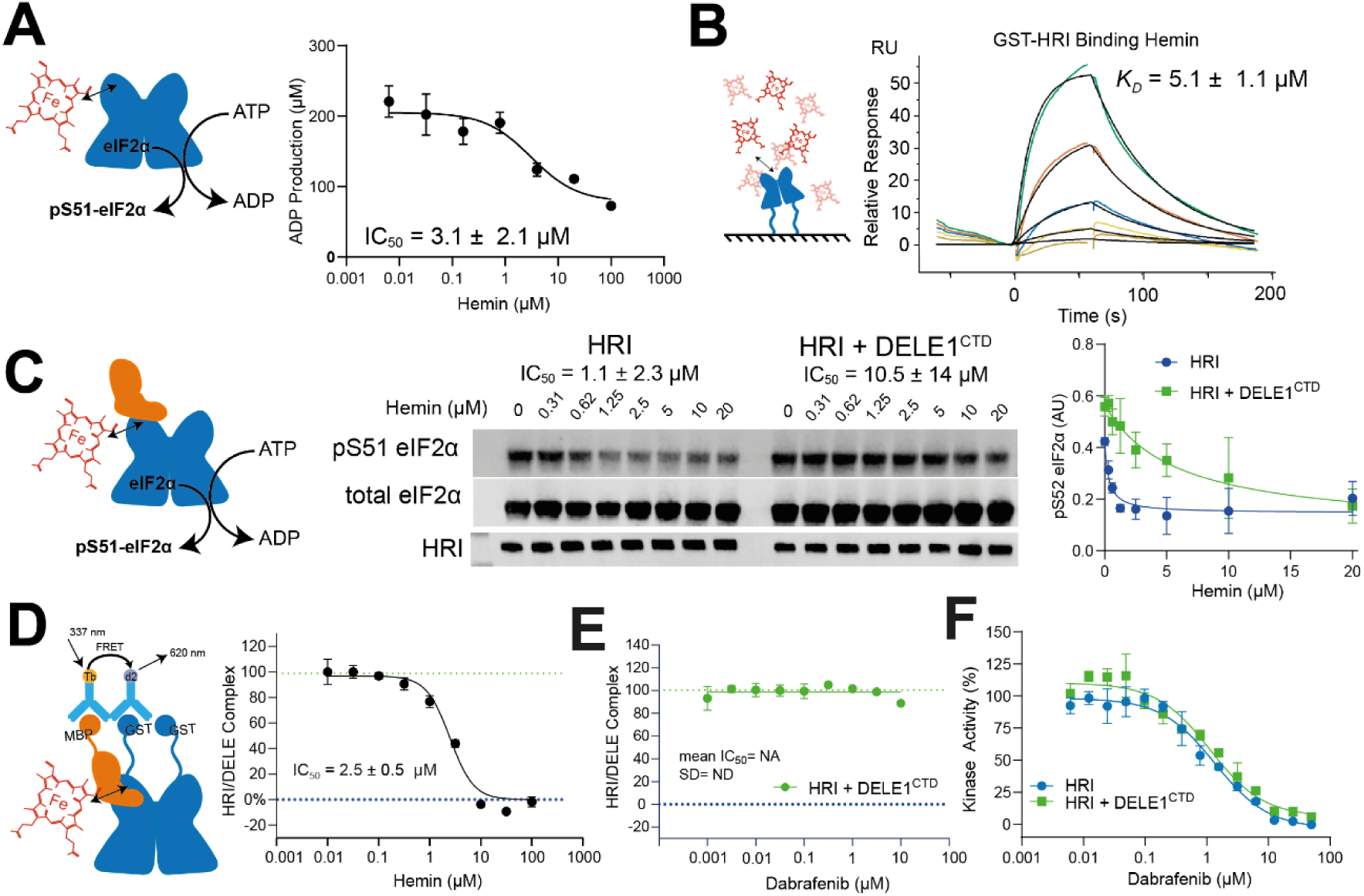
DELE1^CTD^ and Haem competitively bind to HRI to regulate its catalytic activity. (**A**) HRI activity assay with eIF2α substrate, measuring ADP production via an ADP-GLO luminescence assay. Hemin was titrated into the HRI kinase reaction and had a measured IC_50_ of 3.1 µM. Mean and standard deviation of triplicate measurements shown. IC50 value obtained from n=3. (**B**) SPR data showing hemin interaction with immobilised HRI. Representative spectrogram shown. (**C**) DELE1^CTD^ and Haem compete to regulate HRI catalytic activity. Kinase assay measuring product formation (pS51 on eIF2α). Substrate and enzyme concentrations were kept constant with a hemin titration. DELE1^CTD^ reduced HRI sensitivity to hemin approximately 10-fold from 1.1 µM to 10.5 µM. Representative immunoblot shown with total eIF2α and HRI loading controls. Data from three biological repeats for pS51 was quantified and plotted to determine IC_50_ values. (**D**) Hemin binding dissociates DELE1^CTD^: HRI complex formation as measured by TR-FRET. A stable TR-FRET signal formed by complex formation is disrupted by a titration of hemin, with an IC_50_ value of 2.5 µM determined. Mean and standard deviation of triplicate measurements shown. IC_50_ value obtained from n=6. (**E**) Dabrafenib is unable to disrupt the TR-FRET signal associated with DELE1^CTD^: HRI complex formation. Mean and standard deviation of triplicate measurements shown n= 5. (**F**) Inhibition of the HRI/HRI-DELE1^CTD^ kinase activity by Dabrafenib. Mean and standard deviation of triplicate measurements shown n=3.

Since previous studies have primarily investigated the activation of HRI by DELE1^CTD^ in cell-based studies (Yang *et al*, 2023; Bora *et al*, 2025; Sekine *et al*, 2023; Singh *et al*, 2025), we sought to characterise this activation biochemically using purified, recombinant proteins. We incubated 25 nM HRI with equimolar DELE1^CTD^ and 40 µM ATP and measured pS51-eIF2α production following titration of hemin at an 8-point 2-fold serial dilution curve from 20 µM **(Figure 2C, Supplementary Figure 4)**. Strikingly, we observed that the HRI:DELE1^CTD^ complex was far less sensitive to hemin inhibition than HRI alone; HRI had an IC_50_ of 1.1 ± 2.3 µM while HRI:DELE1^CTD^ had an IC_50_ of 10.5 ± 4.5 µM (Welch’s t=test, p = .006). Next, we hypothesised that DELE1^CTD^ may compete with hemin for binding to HRI. To test this, we employed the TR-FRET assay to determine if titration of hemin could perturb the interaction between DELE1^CTD^ and HRI **(Figure 2D).** We found that hemin was able to cause the HRI:DELE1^CTD^ complex to disassociate with an IC_50_ of 2.5 ± 0.5 µM, suggesting a competitive mechanism of displacement.

To determine if competitive displacement of HRI:DELE1^CTD^ was specific to hemin, we next tested a small molecule HRI kinase inhibitor. Previously, we have demonstrated that HRI could be inhibited by the ATP-competitive inhibitor Dabrafenib (Kanta *et al*, 2025). Under the same TR-FRET assay conditions, in contrast to hemin, titration of Dabrafenib had no effect on HRI:DELE1^CTD^ complex formation indicating that occupancy of the kinase domain by Dabrafenib is unable to disrupt complex formation **(Figure 2E)**. We next investigated whether the HRI:DELE1^CTD^ complex could still be inhibited using Dabrafenib. Using the ADP-GLO HRI kinase assay, following titration of Dabrafenib, we found no significant difference between HRI and HRI:DELE1^CTD^ inhibition by Dabrafenib (IC_50_ 1.3 ± 0.45 μM and 1.1 ± 0.55 μM, respectively) **(Figure 2F)**.

### DELE1^CTD^ binds directly to the N-terminal haem binding domain of HRI to occlude haem

We next employed HDX-MS to characterise the structural interaction between HRI and DELE1^CTD^. Previously, we have used HDX-MS to characterise how HRI is allosterically inhibited by hemin binding (Kanta *et al*, 2025). HDX-MS investigates protein structure by measuring the solvent exchange rate of the protein amide backbone with solvent – higher rates of solvent exchange are indicative of unstructured regions of protein and solvent exposure, and decreases of solvent exchange are associated with localised hydrogen bonding or solvent exclusion events at the site of protein interaction (Masson *et al*, 2019; Vinciauskaite & Masson, 2023). By comparing the solvent exchange rate of HRI dimers in the absence and presence of an equimolar concentration of DELE1^CTD^, we observed several decreases in solvent exchange, indicative of a binding event (**Figure 3A**, **Table 1.0)**. Using AlphaFold 3 (Abramson *et al*, 2024), we also created a model of an HRI dimer binding to a single copy of DELE1^CTD^, subsequently imposing a monomer of the previously solved structure of oligomeric DELE1^CTD^ (Yang *et al*, 2023) on the AlphaFold Model (**Figure 3B-C) (Supplementary Figure 5)**.

**Figure 3:**
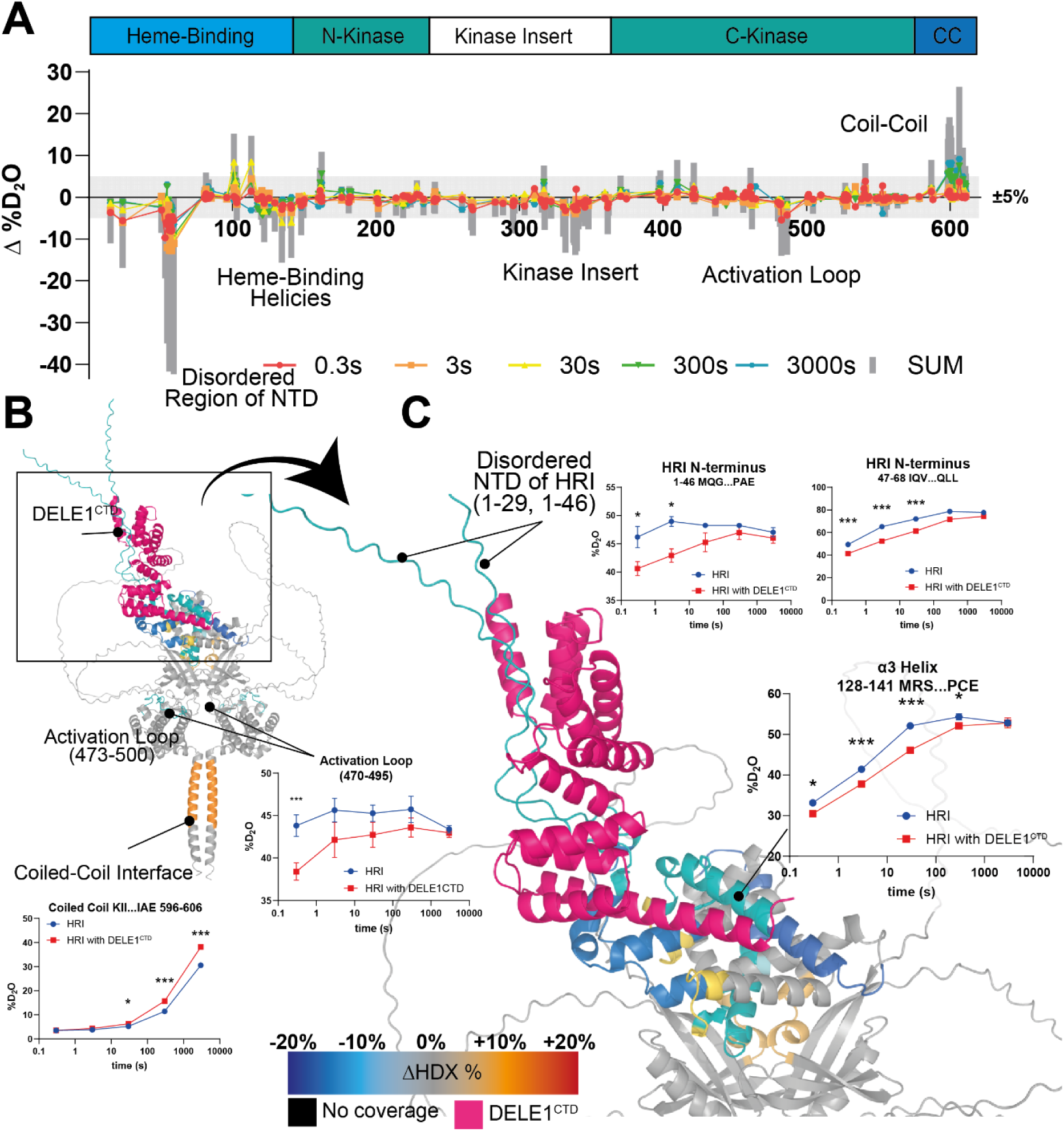
HDX-MS data detailing structural changes in HRI upon complex formation with DELE1^CTD^. (**A**) Overall changes in solvent exchange rate in HRI mapped onto peptides. Increases in solvent exchange rate on **DELE1^CTD^** binding are shown as increase in D_2_O, while decreases are shown as negative values. Each point is a single peptide, with the x-axis being the peptide midpoint (i.e. a peptide spanning residues 5-15 would have a midpoint of 10). The grey bar between +5 and −5% signifies likely significance thresholds. (**B)** Changes in HDX- MS mapped onto an AlphaFold prediction of a complex consisting of 1 DELE1^CTD^ : 2 HRI. DELE1^CTD^ is shown in pink while HRI is coloured according to differences in solvent exchange rate caused by DELE1^CTD^ binding. (**C**) Focus on the interaction site at the haem binding domain of HRI. Each data point in the peptide uptake graphs represents a triplicate measurement with points showing means, error bars standard deviation. Significance p values of non-parametric T tests are indicated by * = 0.05, ** = 0.01, *** = 0.001.

**Table 1.0:** HDX-MS Statistics for HRI DELE1 Binding.

|  | HRI – APO | HRI w/ DELE1<sup>CTD</sup> | DELE1<sup>CTD</sup> | DELE1<sup>CTD</sup> w/<br>HRI |
| --- | --- | --- | --- | --- |
| # of Peptides | 204 | 204 | 138 | 138 |
| Coverage | 94.0% | 94.0% | 97.9% | 97.9% |
| Gaps | 253-254,381-390, 515-521, 615-631 |  | 444-448 |  |
| Av. Length | 17.8 |  | 15.9 |  |
| Redundancy | 8.9 |  | 7.6 |  |
| Time Points | 0.3s<sup>1</sup>, 3s, 30s, 300s, 3000s |  |  |  |
| Controls | Both Non-deuterated and Max Deuteration Controls<sup>4</sup> |  |  |  |
| Replicability<sup>2</sup> | 0.39% |  | 0.67% |  |
| Significance<br>Threshold<sup>3</sup> | > 5%/0.5 Da/Passes t-test |  |  |  |

<sup>1</sup>0.3 s timepoint created by pipetting 3 s by hand with ice-cold solutions.
<sup>2</sup>Mean standard deviation between replicates
<sup>3</sup>A peptide must meet all these criteria to have a significant change in solvent exchange.
<sup>4</sup>Some peptides lack a MaxD assignment. Max Deuteration Controls produced using previously published protocol by Peterle *et al*. (Peterle *et al*, 2022).

Mapping our HDX-MS data onto this model, we found that DELE1^CTD^ primarily interacts with the N-terminus of HRI, especially with the disordered N-terminal region of HRI (residues 1-68) (**Figure 3B-C)**. HDX-MS was able to identify these regions as unstructured due to their very rapid exchange rate (*e.g.* Peptides 1-46, 47-68), and that DELE1^CTD^ binding caused significant decreases in solvent exchange rates (**Figure 3B-C)**. In addition to these changes, we saw significant decreases in the helical bundle of the N-terminus, frequently referred to as the Haem-Binding-Domain (Kanta *et al*, 2025; Ricketts *et al*, 2022; Chen, 2025), including the α3 helix (Residues 108-143 e.g. Peptide 128-141), which accommodates residues H119 and H120, which are critical for haem binding(Cao & Masson, 2026; Igarashi *et al*, 2008). Peptides such as 128-141, which cover the α3 helix show significant decreases, alongside some increases in solvent exchange rates in the α2/α3 hinge (peptides 98-104, 98-106, 100-116), which suggests a remodelling of this domain (**Figure 3B-C)**. In addition to these changes which are close to the predicted interaction site, there were changes in HDX in the kinase insert loop (residues 330-390), and the activation loop (473-500) of the kinase domain. Furthermore, there were also large increases in HDX in the C-terminal coiled-coil (peptide 595-606, 595-613) of HRI which may indicate more distal, allosteric regulation of HRI upon DELE1^CTD^ binding.

The HDX-MS data reveal two main regions of interaction for DELE1^CTD^ binding to HRI (**Figure 4)**. The large N-terminal helix of DELE1^CTD^, the tetratricopeptide repeat (TPR) segment 1 (TPR1), showed moderate decreases in HDX-MS (peptides 225-242, 226-242), and a second helical bundle formed by shorter TPR segments (residues 338-431) also showed significant reductions in HDX (e.g. peptides 338-356, 369-377, 411-429), indicative of the predicted interaction with the disordered N-terminus of HRI (**Figure 4A-B)**.

**Figure 4.**
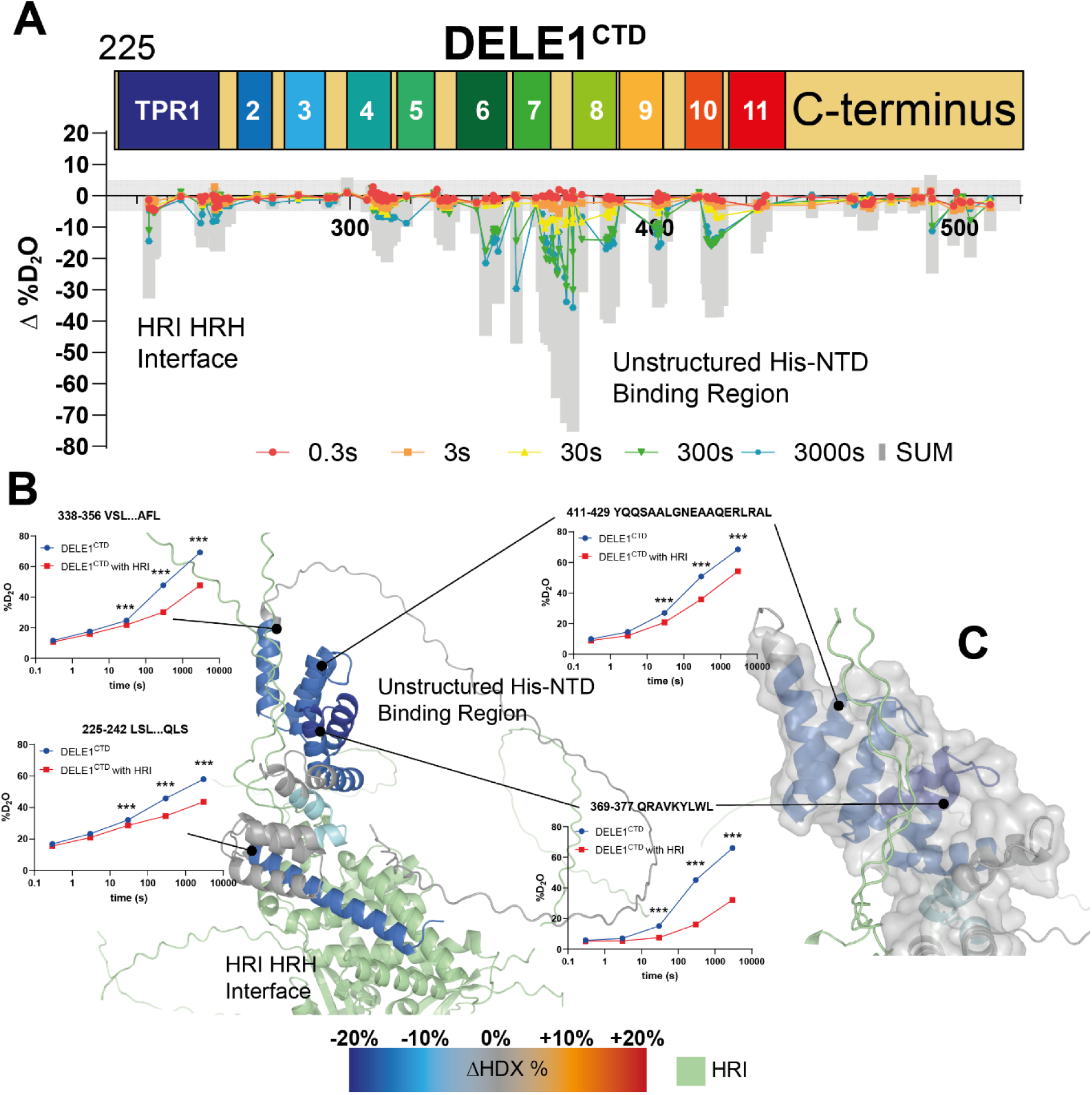
HDX-MS data detailing structural changes in DELE1^CTD^ on complex formation with HRI. (**A**) Overall changes in solvent exchange rate in DELE1^CTD^ mapped onto peptides. Increases in solvent exchange rate on HRI binding are shown as increase in D_2_O, while decreases are shown as negative values. Each point is a single peptide, with the x-axis being the peptide midpoint (i.e. a peptide spanning residues 5-15 would have a midpoint of 10). The grey bar between +5 and −5% signifies likely significance thresholds. (**B)** Changes in HDX-MS mapped onto an AlphaFold prediction of a complex consisting of 1DELE1^CTD^:2 HRI. HRI is shown in green while DELE1^CTD^ is coloured according to differences in solvent exchange rate caused by HRI binding. Each data point in the peptide uptake graphs represents a triplicate measurement with points showing means, error bars standard deviation. Significance p values of non-parametric t-tests are indicated by * = 0.05, ** = 0.01, *** = 0.001.

## Discussion

In this study, we set out to investigate the mechanism by which DELE1^CTD^ binds to and activates HRI. This signalling branch of the integrated stress response (ISR) has been identified as a critical sensor of mitochondrial stress (Guo *et al*, 2020; Fessler *et al*, 2020). In previous research, our group and others have identified an emergent role of the DELE1^CTD^: HRI axis in neurodegenerative disorders such as Parkinson’s disease where it acts as a negative regulator of PINK1-PARKIN mitophagy (Singh *et al*, 2025). Previous studies investigating the interaction between DELE1^CTD^ and HRI have largely been conducted in cell-based co-immunoprecipitation studies with outstanding questions on the mode of binding and quantification of strength of the interaction. In this study we use biochemistry, biophysics and structural mass spectrometry to demonstrate that the DELE1^CTD^ competes with haem for HRI regulation through binding specifically via its TPR1 helix to the N-terminus of HRI.

The structure of the DELE1^CTD^ fragment consisting of residues 225-515, has been solved using cryo-electron microscopy (Yang *et al*, 2023). Yang *et al*. demonstrated that the 225- 515 fragment of DELE1 (DELE1^CTD^) readily oligomerises and proposed that this oligomerisation property is critical to HRI activation – suggesting that HRI may interact with oligomerised DELE1^CTD^ *in cellulo*. However, in our biophysical studies, including mass photometry and SPR **(Figure 1)** we found that it is most likely a single monomer of DELE1^CTD^ (although two cannot be ruled out) rather than an oligomer which interacts with a dimeric HRI. One possible reason for this discrepancy is that Yang *et al*. identified that the TPR1 region of DELE1^CTD^ was part of the oligomerisation interface, and that mutation of TPR1 both prevented oligomerisation and prevented ISR activation. However, using HDX-MS here we have demonstrated that TPR1 plays a role not only in DELE1^CTD^ oligomerisation but also in the HRI interaction (**Figure 4**), suggesting that any mutations that disrupt oligomerisation may also ablate the ability of DELE1^CTD^ to interact with HRI. This is consistent with Fessler *et al*. who demonstrated that the TPR1 region of DELE1^CTD^ was critical for ISR activation via HRI (Fessler *et al*, 2020).

To date it has been unclear whether HRI could bind to both DELE1^CTD^ and haem simultaneously. There have been numerous studies on how haem may inhibit HRI (Cao & Masson, 2026), and these have highlighted that whilst the N-terminal region of HRI is a critical binding site for haem (Kanta *et al*, 2025; Ricketts *et al*, 2022; Igarashi *et al*, 2008), the interaction may be more complex. Haem binding requires HRI dimerization (Ricketts *et al*, 2022), and potential second sites of interaction have been identified within the kinase insertion region (Rafie-Kolpin *et al*, 2000), alongside cysteine and proline residues within the C-lobe of the kinase domain (Igarashi *et al*, 2008). We have also shown previously that haem binding is associated with widespread structural changes in HRI (Kanta *et al*, 2025). It has also been frequently observed in numerous biochemical studies that haem binding produces incomplete inhibition of HRI *in vitro* (Igarashi *et al*, 2011, 2008; Ricketts *et al*, 2022; Kanta *et al*, 2025), suggesting that haem could remain bound alongside any activator of HRI.

In this study we show, using kinase activity assays (**Figure 2C)** and a TR-FRET complex formation assay (**Figure 2D),** that haem binding and DELE1^CTD^ binding are mutually exclusive. Furthermore, we demonstrate using SPR and TR-FRET assays that the affinity of DELE1^CTD^ for HRI is approximately 200-fold greater than the affinity of hemin for HRI (22 nM vs 5 µM, respectively) (**Figures 1 & 2)**, which suggests that even a small amount of DELE1^CTD^ released in the cytosol would be sufficient to compete with and displace the haem bound to HRI when haem levels remain static. We also found that autophosphorylated HRI was still capable of forming a complex with DELE1^CTD^ – suggesting that DELE1^CTD^, after activating HRI, may remain bound during substrate phosphorylation **(Supplementary Figure 2G)**. Furthermore, we observed that while the ATP competitive inhibitor Dabrafenib inhibits HRI both in the absence (as previously reported (Kanta *et al*, 2025)) and presence of DELE1^CTD^, it is incapable of disrupting the HRI/DELE1 ^CTD^ interaction **(Figure 2E&F)**.

The interaction between HRI and DELE1^CTD^ was also investigated using HDX-MS. We have previously used HDX-MS to investigate the wide-spread structural effects of hemin binding to HRI (Kanta *et al*, 2025), and while several regions are conserved between DELE1^CTD^ and hemin, suggesting a common site of interaction, hemin binding produces more extensive changes on the HRI structure than DELE1^CTD^ **(Figure 5A/B)**.

**Figure 5:**
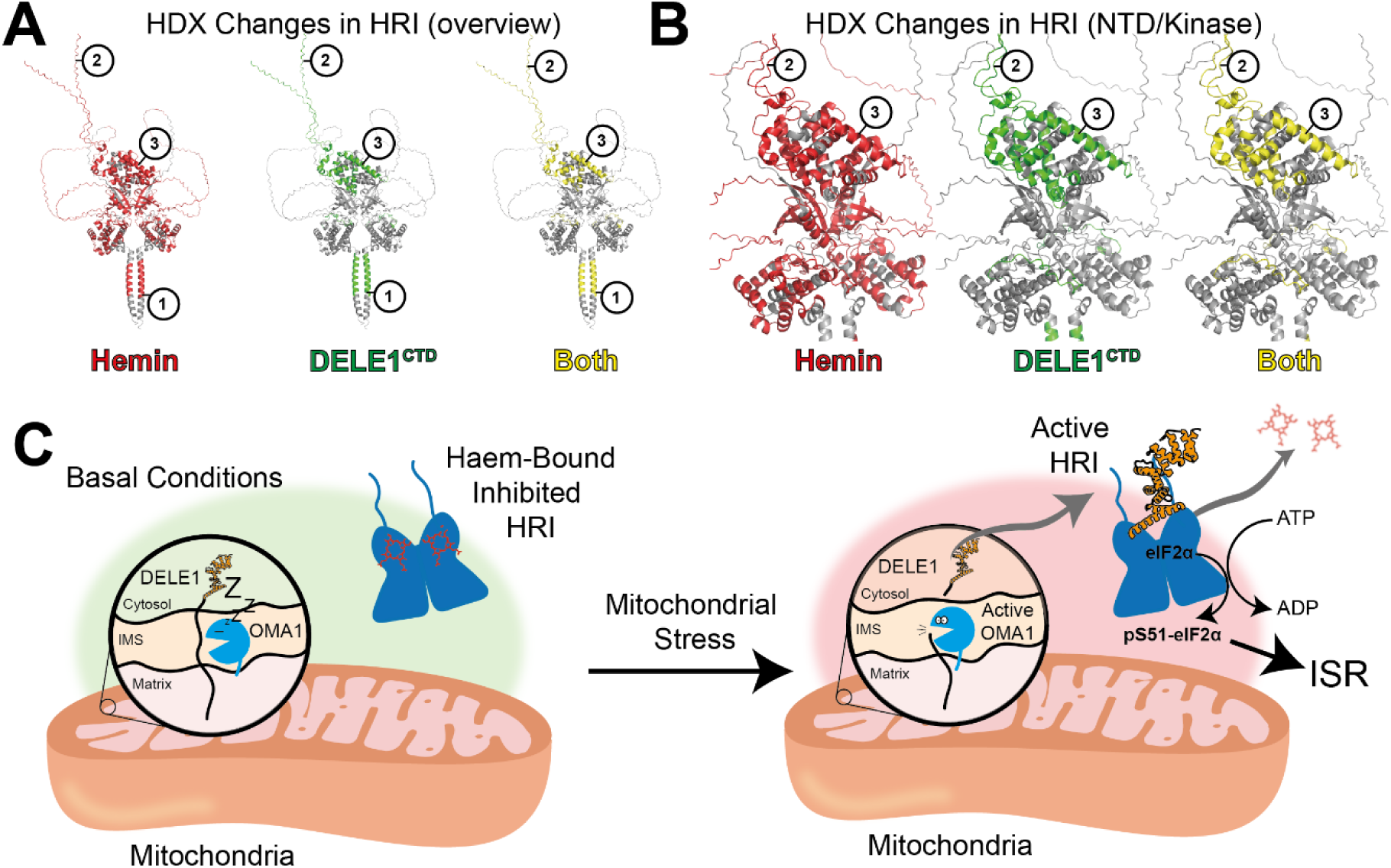
Molecular basis of HRI regulation by Haem and DELE1^CTD^. (**A**) and (**B**) Locations of HRI structural changes on (red) hemin binding, (green) DELE1^CTD^ binding and (yellow) both. Key regions of regulation appear to be (1) the Coil-coil interaction region, (2) the N-terminal disordered region of HRI and (3) the α3 helix of HRI. (**C**) Model of HRI activation by DELE1^CTD^. Under basal conditions, full-length DELE1 resides in the mitochondrial inner membrane, and HRI is bound to haem, autoinhibited in the cytosol. Under mitochondrial stress, the protease OMA1 is activated and cleaves DELE1 to generate a C-terminal DELE1 domain (DELE1^CTD^) that is released into the cytosol. DELE1^CTD^ can then bind to the N-terminus of HRI, displacing haem, and facilitating activation, autophosphorylation and subsequent substrate (eIF2α) phosphorylation, activating the Integrated Stress Response (ISR).

HDX localised the majority of the DELE1^CTD^ interaction interface to the N-terminus of HRI, incorporating both the disordered N-terminal region (residues 1-68) and the haem binding domain of HRI, especially the α3 helix (residues 108-143) **(Figure 3)**. These regions of the N-terminus exhibited reductions in solvent exchange rate, indicative of a site of interaction, but near the α3 helix there were also increases in solvent exchange, which may suggest that there is a structural rearrangement in this domain which results in a decrease in helicity or an increase in solvent exposure. Notably, in the absence of DELE1^CTD^, this unstructured region of HRI appears to extend to the first 70 residues (peptide 50-66 has >50% Deuteration at 0.3 s D_2_O exposure, indicating a lack of secondary structure), and is predicted as unstructured in the AlphaFold prediction. However, on binding of DELE1^CTD^ we see a large decrease in HDX around residues 50-70, which may be due to a combination of both solvent protection from DELE1^CTD^ interaction, and the simultaneous folding of a helix – suggested by AlphaFold modelling which produces a kinked helix for this region of HRI which may then cradle the C-terminus of the TPR1 helix of DELE1^CTD^. Previous studies have made truncations of HRI consisting of the first 160 residues and found that this was sufficient to facilitate co-immunoprecipitation with DELE1^CTD^ (Yang *et al*, 2023), although it was not clear which regions of the N-terminal region specifically were involved in the interaction. More distally on HRI, DELE1^CTD^ also causes other structural changes, notably in the region of the activation loop of HRI, autophosphorylation of which is critical for activation (Rafie-Kolpin *et al*, 2003). The very C-terminus of DELE1^CTD^ is disordered, and we did observe some decreases in solvent exchange (*e.g.* 484-497, 491-505) – there is potential that these more distal changes from the α3/TPR1 interaction site could be mediated by this C-terminus, but this will require further experimentation.

We also observed that the C-terminal coiled-coil region of HRI undergoes very significant increases in HDX on DELE1^CTD^ binding. Our previous HDX-MS study on HRI showed that both Dabrafenib and hemin binding caused changes in this coil-coil interaction region, suggesting that this structural element appears to be quite sensitive to HRI activity, despite its apparent distance from either the hemin binding site or the kinase domain active site.

The DELE1^CTD^ binding interface falls into two key areas; the long TPR1 helix (residues 225-260), which AlphaFold positions as parallel to the HRI α3 helix, and then a second bundle of TPR helices (TPRs 6-11) which appear to interact with the unstructured N-terminus of HRI **(See Figure 4**). Previous research by Fessler *et al*. had demonstrated that deletion of TPR1 blunted the activation of the ISR but still maintained an interaction suggesting that TPR1/ α3 helix interaction is not necessary for interaction, but critical for activation of HRI, with our HDX data pointing towards a role of the TPR1 preventing haem inhibition of HRI. The second interaction site, the TPR bundle consisting of residues ∼330 to 415, appears to wrap around the N-terminal disordered segments of HRI, clamping the DELE1^CTD^ in place.

Whilst our manuscript was in preparation, highly complementary findings were published by Zhang *et al*.(Zhang *et al*, 2026). Using mammalian cell-based studies alongside pull-down assays, gel mobility shift analysis and protein cross-linking, Zhang *et al*. also demonstrated that each HRI dimer likely interacts with a single monomer or perhaps a dimer of S-DELE1, and that this interaction likely occurs via the α3 helix of HRI. They also found through spectroscopic means that there is mutual exclusivity between DELE1 binding and haem binding to HRI, and haem binding/release was also associated with structural changes distal to the N-terminus of HRI, including in the CTD and the activation loop of HRI.

In conclusion, we present a structural and biophysical model for how DELE1 ^CTD^ and haem binding compete for regulation of HRI activity (**See Figure 5C)**. This expands our mechanistic understanding of the crucial role this interaction plays in sensing mitochondrial stress and its emerging relevance to an increasing number of diseases.

## Materials and Methods

### Materials

| Reagent/Resource | Manufacturer/Source | Identifier/Catalogue Number |
| --- | --- | --- |
| **A) Antibodies** |  |  |
| Anti-HRI antibody | Invitrogen | 7H3L3 |
| Anti-Total eIF2α antibody | Santa Cruz Biotechnology | SC-133132 |
| Anti-pSer51 eIF2α antibody | Cell Signaling Technology | 9721S |
| IRDye680 Goat anti-Mouse antibody | LI-COR | 926-68070 |
| IRDye 800CW Donkey anti-Rabbit antibody | LI-COR | 926-32213 |
| Anti-Histidine antibody (His Capture Kit) | Cytiva | 28995056 |
| Anti-GST antibody (GST Capture Kit) | Cytiva | BR100223 |
| HTRF Anti-MBP mAb Tb-Conjugate (donor antibody) | Revvity | 61MBPTAB |
| HTRF Anti-GST mAb d2-Conjugate (acceptor antibody) | Revvity | 61GSTDLB |
| **B) Chemicals, Enzymes and Other Reagents** |  |  |
| BL21 Star (DE3) E. coli competent cells | New England Biolabs | C2527 |
| Isopropyl β-D-1-thiogalactopyranoside (IPTG) | FORMEDIUM | IPTG025 |
| Benzonase | Novagen | 70664 |
| cOmplete EDTA-free Protease Inhibitor tablet | Roche | 04693132001 |
| Lysozyme | Sigma-Aldrich | L6876 |
| DNase I | Sigma | 04536282001 |
| HisTrap HP Column, 5 mL | Cytiva | 17524801 |
| Q HP Column, 5 mL | Cytiva | 17115401 |
| VIVASPIN 10k MWCO Concentrator | Sartorius | VS2002 |
| Lambda Protein Phosphatase (Lambda PP) | New England Biolabs | P0753S |
| Superdex 200i 10/300 GL column | Cytiva | 28990944 |
| Gel Filtration HMW Calibration Kit | Cytiva | 28403842 |
| Superose 6 Increase 10/300 GL column | Cytiva | 29091596 |
| Ni-NTA beads | Thermo | R90101 |
| Glutathione Agarose Beads | Thermo | 16100 |
| Hemin Chloride | Merck Millipore | 3741 |
| Dabrafenib (10 mM in 100% DMSO) | Med Chem Express | HY-14660 |
| HTRF PPI Terbium Detection Buffer | Revvity | 61DB10RDF |
| 1536-well polystyrene F-bottom HiBase white plate (HTRF) | Greiner | 782075 |
| ADP-Glo Kinase Assay kit | Promega | V6930 |
| 384- Well White, PS, F-bottom plate (ADP-GLO) | Greiner | 781904 |
| Sensor Chip CM5 | Cytiva | BR100530 |
| Bovine Serum Albumin | Merck | A7906100G |
| Culture Well Reusable Gasket (50-3 mm DIA x 1 mm Depth) | Grace Bio-Labs | GBL103250 |
| Enzymate BEH Pepsin Column | Waters | 186007233 |
| ACQUITY UPLC BEH C18 VanGuard Pre-Column | Waters | 186003975 |
| ACQUITY UPLC BEH C18 Column, 1.7 μm, 1.0 x 100 mm | Waters | 186002346 |
| PHERASTAR FSX plate reader | BMG LABTECH |  |
| BIAcore 8K+ instrument | Cytiva |  |
| Refeyn TwoMP mass photometer | Refeyn |  |
| Odyssey FC Imaging System | LI-COR |  |
| Select Series cIMS mass spectrometer | Waters |  |
| iBlot 2 dry blotting system | Invitrogen |  |
| **C) Software** |  |  |
| DiscoverMP v2.5 | Refeyn |  |
| Protein Lynx Global Server | Waters |  |
| HDExaminer 3.3 | Trajan Scientific |  |
| Image Studio | LI-COR |  |
| GraphPad Prism 10 | GraphPad |  |
| Biacore Insight Evaluation v6.0.7.1750 | Cytiva |  |

### Methods

#### Molecular Biology and Cloning

Human HRI cDNA was subcloned into a pOPTH plasmid with an N-terminal 6His TEV-cleavable tag or an N-terminal GST-tag. A full list of plasmids used in this manuscript is listed in Supplementary Table 1. All constructs were verified by The Sequencing Services (School of Life Sciences, University of Dundee) and are now available to request via the Medical Research Council Protein Phosphorylation and Ubiquitylation Unit (MRC PPU) Reagents and Services website (https://mrcppureagents.dundee.ac.uk/).

#### HRI and DELE1 protein Expression and Purification

##### His-Tag

Protein was expressed in BL21 (DE3) (NEB) *E. coli* cells. Cells were grown at 37 °C in 2xTY media until an optical density of 0.7 at 600 nm was achieved and temperature was lowered to 18 °C, where protein expression was induced on the addition of 0.8 mM IPTG. Cells were left to express protein for 16 h at 18 °C. A pellet generated from 3 L of media was lysed in 100 mL Lysis Buffer (20 mM Tris pH 8.0, 500 mM NaCl, 20 mM Imidazole, 2 mM beta-mercaptoethanol (BME), with 1 µL benzonase, (Novagen) and 2 EDTA-free Protease inhibitor tablet (Roche)) via probe sonication. Lysate was centrifuged at 4 °C at 40,000 *g* for 45 minutes. The supernatant was filtered through a 0.45 µm filter and then loaded onto a 5 mL HisTrap HP Column (Cytiva) equilibrated in Lysis Buffer. After washing with 10xCV with lysis buffer and a mAU of <100 achieved, a gradient of NiNTA Buffer B (20 mM Tris pH 8.0, 500 mM NaCl, 200 mM Imidazole pH 8.0, 2 mM BME) was applied. Fractions were analysed using SDS-PAGE to determine HRI-containing fractions.

HRI containing fractions were then diluted 1:1 with Q_O_ Buffer (20 mM Tris pH 8.0, 2 mM BME), and then passed over a 5 mL Q HP (Cytiva) column, equilibrated in Q_A_ Buffer (20 mM Tris pH 8.0, 50 mM NaCl, 2 mM BME), at a rate of 2 ml/min. A gradient of increasing Q_B_ buffer (20 mM Tris pH 8.0, 1 M NaCl, 2 mM BME) was then initiated, with HRI eluting at approximately 600 mM NaCl. HRI containing fractions were then concentrated using a VIVASPIN 10k MWCO Concentrator (Sartorius) until a volume of <1 mL was produced. HRI was then dephosphorylated using Lambda Protein Phosphatase (Lambda PP (New England Biolabs)), with 800 units of phosphatase, and the reaction left at 4 °C for 16 hours. Finally, dephosphorylated HRI was injected onto a Superdex 200i GL 10/300 (Cytiva) equilibrated in Gel Filtration Buffer (20 mM HEPES pH 7.5, 150 mM NaCl, 2 mM TCEP). HRI containing fractions, eluting at ∼10.5 mL, were pooled and concentrated with a VIVASPIN 10k MWCO Concentrator (Sartorius) until a concentration of 1.5 mg/mL was achieved. HRI was then frozen in liquid nitrogen at stored at −70 °C. Gel Filtration calibration was conducted using Cytiva Gel Filtration HMW Calibration Kit (28403842) with the manufacturer’s suggested mix of proteins for the Superdex 200i GL 10/300 column, using the Gel Filtration Buffer as described above.

##### GST-Tag

Expressed and lysed as His-tagged HRI, with the soluble fraction which was incubated with pre-equilibrated GST beads (4 mL) in a Duran bottle (rolling, 2 h, 4 °C). The mixture was applied to a gravity column and then washed with 150 mL wash buffer (20 mM Tris-HCl (pH 8.0), 500 mM NaCl, 2 mM TCEP). GST-HRI was then eluted using elution buffer (20 mM Tris-HCl (pH 7.5), 500 mM NaCl, 50 mM reduced glutathione, 2 mM TCEP). Anion-exchange, dephosphorylation and gel filtration were the same as described for His-HRI.

##### DELE1 Purification

Both Full-length and DELE1^CTD^ were purified as previously reported – S- DELE1 was purified the same as His-MBP-DELE1^CTD^ (Yang *et al*, 2023). For His-MBP- DELE1^CTD^, BL21(DE3) *E.coli* were grown at 37 °C using auto-induction media until an O.D of 0.8, before a drop of temperature to 17 °C and allowed to continue growing overnight or induced with 1 mM IPTG for 16h. Cells were resuspended in lysis buffer (25 mM Tris-HCl (pH 8.0), 500 mM NaCl, 10% glycerol, 1 mM TCEP containing EDTA Free-protease inhibitor cocktail (Roche), lysozyme at 0.1 mg/mL, 20 mM imidazole and 10 μg/mL DNase1) and then lysed cells using a sonicator (10sec on; 10sec off pulse for a 4 min cycle), followed by ultra-centrifugation at 53,343 x *g* for 40 min. Supernatant was incubated with 3 ml NTA-beads in a gravity column for 2h on a roller at 4 °C. Beads were washed with 100 mL wash buffer (25 mM Tris-HCl (pH 8.0), 500 mM NaCl, 10% glycerol, 1 mM TCEP and 20 mM imidazole), followed by elution with 50 mL Elution Buffer (25 mM Tris-HCl (pH 8.0), 500 mM NaCl, 10% glycerol, 1 mM TCEP and 200 mM imidazole). Eluted protein was confirmed by SDS-PAGE then concentrated and purified further by SEC on a Superose 6 increase 10/300 GL column previously equilibrated with gel filtration buffer (25 mM HEPES (pH 7.5), 150 mM NaCl, and 1 mM TCEP). DELE1^CTD^ containing fractions were pooled and concentrated to ∼15 mg/mL with an average yield 3.8 mg/L of culture. For Full-Length DELE1 a similar protocol was used with an alternative lysis buffer (25 mM Tris-HCl (pH 8.0), 500 mM NaCl, 10% glycerol, 1 mM TCEP, 20 mM imidazole, 0.5 % DDM, 0.1 mg/mL lysozyme, protease inhibitor cocktail (Roche)) and wash buffer (25 mM Tris-HCl (pH 8.0), 500 mM NaCl, 10% glycerol, 0.1% DDM, 20 mM imidazole and 1 mM TCEP) and the gel filtration buffer (25 mM HEPES, 150 mM NaCl, 0.1% DDM and 1 mM TCEP). Yield 1.2mg/L of culture.

##### Calibrated Gel Filtration

Gel Filtrations Standards (Gel Filtration Standard BioRad #1511901 for Superose 6 increase 10/300 GL, Cytiva Gel Filtration Calibration Kit # 28403842 Superdex 200i 10/300 GL) were run to the manufacturer stipulations with a 0.5 mL loop at a flow rate of 0.5 mL/min in the protein’s gel filtration buffer.

##### eIF2α Expression and Purification

Expression and purification of recombinant human eIF2α was conducted as described previously (Inglis *et al*, 2019). DNA encoding full-length human eIF2α (NCBI reference number: NP_004085.1) was inserted into the vector pOPTH with an N-terminal His6 tag followed by a TEV protease site. The plasmid was transformed into chemically competent BL21 Star (DE3) cells, and cells were grown overnight before being inoculated to a 50 mL starter culture in 2xTY media containing 0.1 mg/mL Ampicillin. The starter culture was incubated at 37 °C for 90 minutes, then 10 mL starter culture was added to 4 x 900 mL 2xTY media containing Ampicillin. Cultures were incubated at 37 °C until the optical density reached 0.7, and then protein expression was induced by the addition of 0.3 mM isopropyl β-D-1- thiogalactopyranoside (IPTG). Cells were grown for a further 3 hours at 37 °C before being harvested, washed with ice-cold phosphate-buffered saline and frozen in liquid nitrogen. Bacterial cell pellets were lysed in 100 mL Lysis Buffer (20 mM Tris-HCl pH 8.0, 100 mM NaCl, 5 % v/v glycerol, 2 mM BME 0.5 mg/mL Lysozyme (Sigma L6876), 2 U/mL Benzonase, one cOmplete EDTA-free protease inhibitor tablet (Roche 04693132001) per 50 mL of Buffer). Cells were lysed using a probe sonicator for 5 minutes (10 s on/ 10 s off) and then centrifuged at 140,000 *g* for 45 min at 4 °C. The supernatant was filtered through a 0.2 µm syringe filter before being loaded onto a 5 mL HisTrap HP Column (Cytiva 17524801) equilibrated in low imidazole Buffer A (20 mM Tris pH 8.0, 100 mM NaCl, 5 % v/v glycerol, 10 mM imidazole pH 8.0, 2 mM BME), followed by the elution of protein via a gradient using Buffer B (20 mM Tris pH 8.0, 100 mM NaCl, 5% v/v glycerol, 200 mM Imidazole pH 8.0, 2 mM BME). Protein gel filtration purification then proceeded as described for HRI. Proteins were concentrated to ∼10 mg/mL and then snap frozen in liquid nitrogen.

##### Hemin Solution

For SPR and TR-FRET, 1 mM Hemin stocks were created via addition of 100% DMSO (this was to allow for direct comparison with Dabrafenib (also in 100% DMSO)). For ADP-GLO, Western blot assays, a 5 mM stock was produced using 32.6 mg hemin chloride which was then dissolved in 1000 μL 1 M NaOH and 8.0 ml ethylene glycol. The pH was adjusted using 1 M HCl to pH 7.5 solution, and the volume adjusted to 10 mL with the addition of ultrapure water. The hemin solution was stored at −20 °C.

##### TR-FRET Assay

3 nM GST-HRI and 5 nM His-MBP-DELE1^CTD^ were pre-incubated for 15 min at 25°C in a volume of 2 µL in HTRF PPI Terbium Detection Buffer (Revvity, 61DB10RDF)) in a 1536 well, Polystyrene, F-bottom, hibase, white plate (Greiner, 782075). This solution was then mixed with 2 µL of a donor antibody (HTRF Anti-MBP mAb Tb-Conjugate (Revvity, 61MBPTAB) /acceptor (HTRF Anti-GST mAb d2-Conjugate (Revvity, 61GSTDLB)) antibody mix in PPI Terbium Detection Buffer. The 4 µL reaction was then incubated for 2 h at 25°C while shielded from light. Fluorescence signals were then measured using a PHERAstar FSX plate reader in HTRF mode with simultaneous measurements of both 620 nm cryptate and 665 nm acceptor emissions, with an integration delay (lag time): 60 µs, integration time: 400 µs, with the number of flashes: 27. For compounds/protein dosing, 10 nL of compound/protein was dispensed using Echo650 into the plate prior to addition of GST-HRI/His-MBP-DELE1^CTD^ to the plate. Data analysis: HTRF ratio was calculated using the following equation: ((HTRF at 665 nm / HTRF at 620 nm) x 10000). To obtain a K_D_ for DELE1^CTD^ an 8-point titration of unlabelled His-HRI (300, 100, 33.3, 11.1, 3.7, 1.2, 0.4, 0.1 nM) was used to disrupt the TR-FRET signal.

##### SPR Assays

Surface Plasmon Resonance (SPR) data were collected using a BIAcore 8K+ instrument (Cytiva) by CROs Selvita and Hitgen. Both His-tagged and GST-tagged HRI were used in this analysis. For His-tagged HRI, an anti-histidine antibody was covalently coupled to a Sensor Chip CM5 using 1-ethyl-3-(3-dimethylaminopropyl)-carbodiimide hydrochloride (EDC) and N-hydroxysuccinimide (NHS) activation and ethanolamine deactivation as per the manufacturer’s instructions (Cytiva His Capture Kit, 28995056). A similar process for GST-tagged HRI was conducted using the GST Capture Kit (Cytiva, BR100223). 100 µg/mL HRI was then captured on the chip to an RU of ∼850 (Hemin binding) or 130 for DELE1^CTD^ binding. Multi-cycle kinetics titration modes were then employed with 60 second association time, followed by 120 sec dissociation steps, at a flow rate of 30 µL/min. The assay buffer was HBSP (0.01 M HEPES pH 7.4, 0.15 M NaCl, 0.005% v/v Surfactant P20).Hemin Interaction specifics: A 10-point concentration curve (30, 10, 3.3, 3, 1.1, 1, 0.37, 0.3, 0.1, 0.04 µM) of Hemin in 1% DMSO was analysed against and immobilised GST-HRI and His-HRI. GST-HRI and HIS-MBP-DELE1^CTD^: A 6-point concentration curve was prepared (600, 200, 67, 22, 7.4, and 2.5 nM) of His-MBP-DELE1^CTD^. In parallel, a reference channel with only the antibody (i.e. no HRI) was used to check for non-specific binding. All data were doubly referenced by subtraction for both the reference channel and from injections of the buffer alone. Data were analysed using Biacore Insight Evaluation v6.0.7.1750 (Cytiva). Final kinetics data reported (K_D_ etc.) are averages of at least three independent experiments with both tags (*i.e.* both His and GST approaches), with individual sensograms in figures as examples.

##### pS51-eIF2alpha kinase assays

Concentrations of HRI, eIF2α, DELE1^CTD^ were used as described in the results/ figure legends (typically 25 nM HRI, 5 μM eIF2α, 25 nM DELE1^CTD^) with 40 μM ATP (PROMEGA). Reactions were conducted in kinase assay buffer (20 mM HEPES pH 7.5, 150 mM NaCl, 5 mM MgCl_2_, 1 mM TCEP) for 20 mins at room temperature. Assays were quenched via the addition of SDS-Loading buffer and 2 minutes of boiling at 95 °C.

##### Western immunoblots

Gel transfers from SDS-PAGE gels onto nitrocellulose membranes were conducted using a semi-dry method with the iBlot 2 P0 method. All blocking buffers were 4% BSA in TBST. Antibodies used: HRI (Invitrogen 7H3L3, Rabbit, 1:500), Total eIF2α (Santa Cruz Biotechnology, SC-133132, 1:500), pSer51 eIF2α (Cell Signalling Technology 9721S, 1:250), IRDye680 Goat anti Mouse (Licor 926-68070), IRDye 800CW Donkey Anti Rabbit (Licor 926-32213). All images taken using Licor Odyssey FC system and images analysed using ImageStudio. Immunoblots representative of a minimum of three repeats. Quantified immunoblots were further analysed using GraphPad prism software, with IC_50_ values being determined using a standard non-linear regression analysis. Each experiment conducted a minimum of three repeats, not including technical repeats (*i.e.* re-blotting of the same kinase assay).

##### ADP-GLO Kinase Assays

Using a PROMEGA ADP-GLO Kinase assay, 50 nM HRI was incubated with 2 μM eIF2α and 250 μM ATP at the stated concentrations in Kinase Reaction Buffer (20 mM HEPES pH 7.5, 150 mM NaCl, 5 mM MgCl_2_, 1 mM TCEP) for 45 min at room temperature, until the reaction was quenched using the ADP-GLO Reagent. 4 µL kinase reactions were conducted in Greiner 384 well plate (781904) before quenching and development using the ADP-GLO/Kinase Detection Reagent as per the manufacturer’s instructions. An ADP/ATP curve was created using the kit for calculation of relative ADP concentrations after kinase reaction quenches. Luminescence measured using a Pherastar (BMG LABTECH). Data analysis conducted using Graphpad Prism 10 (Graphpad). A simple linear regression model for the ADP calibration curve was used to determine the relationship between luminescence and ADP concentration. Non-linear regression analysis (typically [Inhibitor] vs. response (three parameters) was used in Hemin experiments.

##### AlphaFold Prediction

The human full-length HRI (1-630) DELE1^CTD^ (225-515) sequences were used as a for the structure prediction using for AlphaFold 3(Abramson *et al*, 2024).

##### Mass Photometry

Mass determination of solution phase HRI and DELE1^CTD^ were conducted as previously described(Kanta *et al*, 2025). Data were collected using a Refeyn TwoMP instrument. Experimental data were collected using Grace Bio-Labs Culture Well Reusable 50-3mm DIA x 1mm Depth Gaskets and High Precision glass microscope slides, repeatedly cleaned in ultrapure water and isopropanol and dried using a stream of nitrogen gas. Data were obtained through the collection of mass photometry videos. Prior to HRI/DELE1 data collection, mass calibration was conducted using BSA (66 kDa) and aldolase (160 kDa). By fitting Gaussian functions to ratiometric contrast values obtained from the protein standards using the DiscoverMP v2.5 software (Refeyn), a linear mass calibration was obtained. HRI proteins were diluted to 100 nM in 20 mM HEPES pH 7.5, 150 mM NaCl, and 2 mM TCEP, placed on the slide, and then a one-minute mass photometry video was collected. Data was analysed using the DiscoverMP v2.5 software. Data presented are representative histograms of one of three repeats.

##### HDX-MS Sample Preparation

7 µM HRI was incubated with or without 7 µM His-MBP- DELE1^CTD^ for 30 minutes in Protein Dilution Buffer (25 mM HEPES pH 7.5, 150 mM NaCl, 2 mM DTT). 5 µL of this sample was diluted with 45 µL of Deuteration Buffer (25 mM HEPES pH 7.5, 150 mM NaCl, 2 mM DTT, 94.4 % D_2_O) (final D2O concentration = 85 %) for timepoints 3/ 30/ 300/ 3000 s, before being quenched with 20 µL of ice-cold Quench Solution (6 M Urea, 2% Formic Acid), and being snap frozen in liquid nitrogen and stored at – 70 °C. A further timepoint, 0.3 s, was achieved by incubating both the sample and deuteration buffer on ice and then conducting a 3 s exchange reaction. Each exchange reaction was conducted independently three times. Max deuteration controls were produced also using the maxD protocol (Peterle *et al*, 2022).

##### HDX-MS Data Acquisition

Data acquisition was conducted broadly as previously described (Kanta *et al*, 2025). Samples were rapidly thawed at room temperature and then injected into automated HDX-MS fluidics and UPLC manager system (Waters). Samples were loaded into a 50 µL loop and then subsequently digested using a Waters Enzymate BEH Pepsin Column (Part No. 186007233) in a 0.1 % Formic Acid solution with a flow rate of 200 µL/min at 20 °C. Peptic peptides then flowed onto a Waters ACQUITY UPLC BEH C18 VanGuard Pre-Column at 1 °C (Part No. 186003975). After digestion, the flowpath was changed to elute the peptides via a Waters ACQUITY UPLC BEH C18 1.7 µm 1.0 x 100 mm reverse phase column (Part No. 186002346). A gradient from 0-85% 0.1% Formic Acid/ Acetonitrile, conducted at 1 °C with a 40 µL/min flowrate was used to elute deuterated peptides which were then ionised using an ESI source. Mass spectrometry data were collected using a Waters Select Series cIMS instrument, from a 50-2,000 m/z range with the instrument in HDMSe mode. A single pass of the cyclic ion mobility separator (with a cycle time of 47 ms) was conducted. A blank sample of protein dilution buffer with quench was run between samples, and carryover was routinely checked to be <1% intensity of the prior sample.

##### HDX-MS Data Analysis

Peptide sequence identification was conducted using Protein Lynx Global Server (Waters). Minimum inclusion criteria were a minimum intensity of 5000 counts, minimum sequence length 5, maximum sequence length 35, a minimum of 3 fragment ions, a minimum of 0.1 products per amino acid, a minimum score of 5.0, a maximum MH+ Error of 10 ppm. Subsequent analysis and determination of deuteration values conducted using HDExaminer 3 (Trajan Scientific). Experimental design, data acquisition, analysis, and reporting are in line with the community agreed recommendations(Masson *et al*, 2019).

## Supporting information

DELE1_HDX

HRI_HDX

## Acknowledgements

We thank the MRC PPU Reagents and Service group for help in protein production and plasmid construction. GRM thanks Dr Anna Plechanovova and Prof. Ron Hay for help with Mass Photometry. SPR (Selvita and Hitgen) and TR-FRET (Selvita) assays were conducted by Selvita and HitGen Inc. as a contract research organisation. Aspects of this work were funded by The Royal Society, research grant: RGS\R1\231147. HDX-MS equipment was funded by BBSRC Capital Equipment Fund BB/V019635/1. VV is supported by an MRC iCase Studentship (grant number MR/R01579/1). This work was supported by SPARK NS and the Medical Research Council (UKRI2546). M.M.K.M. is supported by Parkinson’s UK and the UK Dementia Research Institute through UK DRI Ltd, principally funded by the Medical Research Council. MC is supported by a China Scholarship Council (CSC) PhD studentship.

## CRediT Author Contribution

**Olawale Raimi:** Conceptualisation, Resources, Investigation **Min Cao: r**esources, Investigation **Shannon Richardson:** Resources, Investigation **Rupam Bhattacharjee:** Resources, Investigation **Andrew C. H. Liu:** Resources, Investigation **Vanesa Vinciauskaite**: Resources **Carine De Marcos Lousa:** Conceptualisation, Resources, Supervision, Investigation, Funding acquisition, Writing — review and editing **Miratul M.K. Muqit:** Conceptualisation, Resources, Supervision, Investigation, Funding acquisition, Writing — review and editing. **Glenn R Masson**: Conceptualisation, Resources, Supervision, Investigation, Funding acquisition, Methodology, Writing — original draft, Project administration, Writing — review and editing.

## Data Availability Statement

All data and reagents are available from the authors upon request. Uncropped western immunoblots are included as Supplementary Data. HDX-MS Data available on PRIDE Deposition PXD083724.

## Competing Interests

M.M.K.M. is a member of the Scientific Advisory Board of Montara Therapeutics Inc. and a scientific consultant to Mission Therapeutics. The other authors declare that they have no competing interests.

---

**Supplementary Figure 1:**
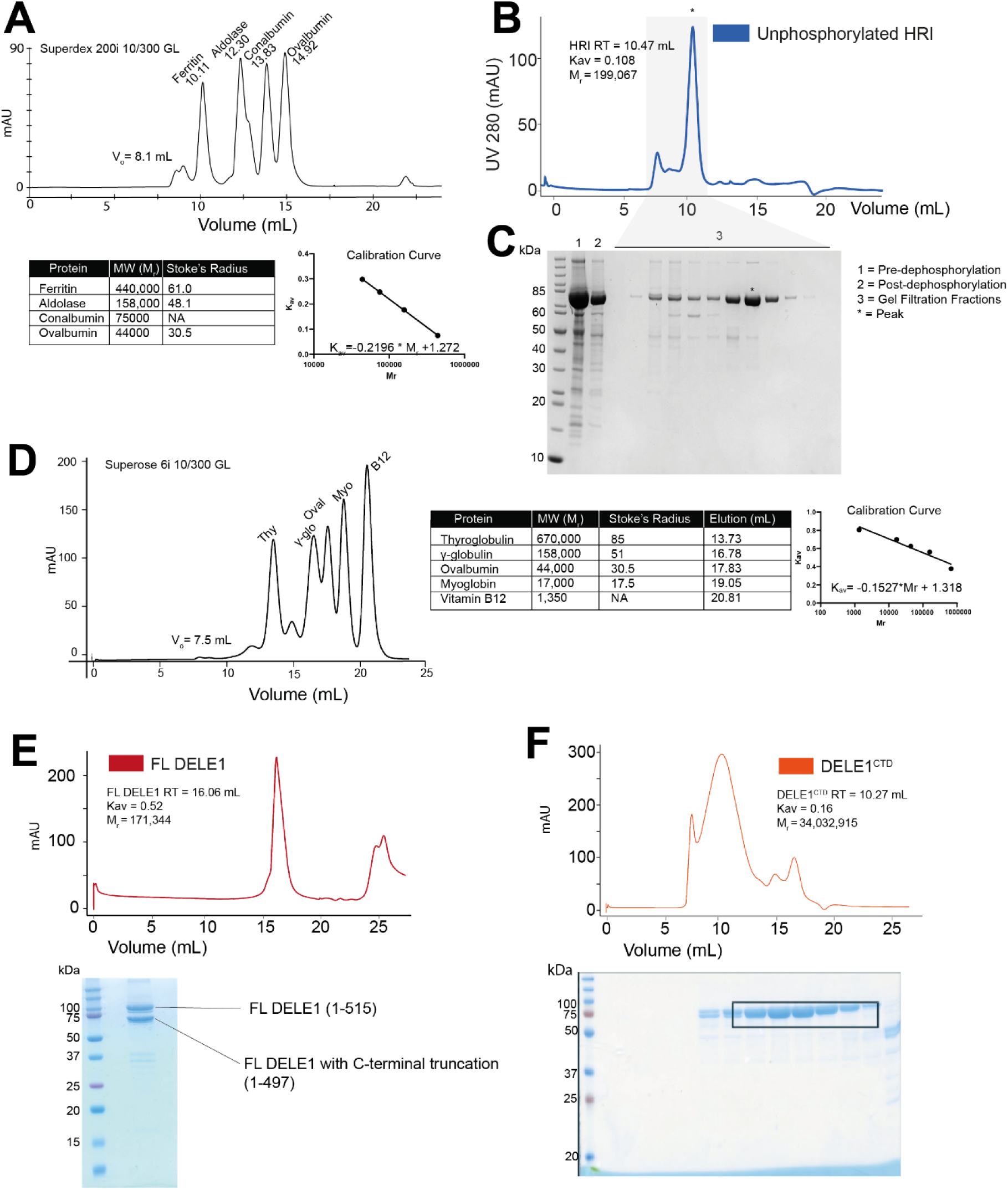
Protein Purification of HRI, DELE1-FL and DELE1^CTD^. (**A**) Calibrated Size Exclusion Chromatography (SEC) Details for Superdex 200i 10/300 GL column. (**B**) SEC of dephosphorylated His-TEV-HRI (**C**) SDS-PAGE of SEC fractions of HRI. (**D**) Calibrated SEC of Superose 6 increase 10/300 GL. (**E**) Gel filtration of Full Length DELE1. A dual band was observed on SDS-PAGE; mass spectrometry analysis showed this was a C-terminal truncation of FL DELE1. (**F**) SEC profile of DELE1^CTD^ and SDS-PAGE.

**Supplementary Figure 2:**
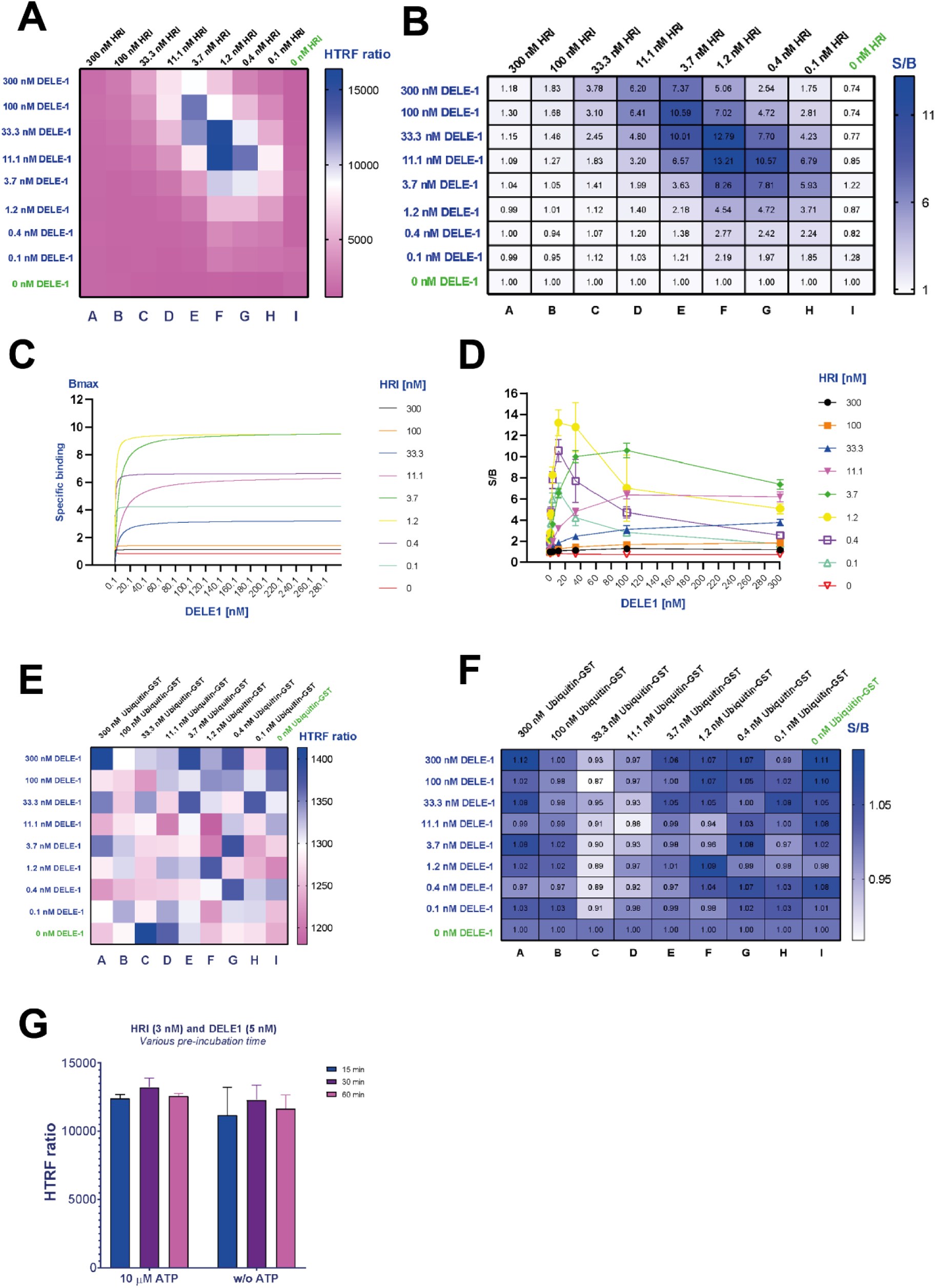
TR-FRET Optimisation. **(A)** 2-way titration of DELE1^CTD^ and HRI. HTRF = Homogeneous time-resolved fluorescence. (**B**) 2-way titration showing Signal/Background ratio. (**C/D**) Curves and Datapoints showing saturation of fluorescence signal (and eventual hook effect). (**E/F**) Negative control protein GST-Ubiquitin titrated against DELE-1. (**G**) Pre-incubation of HRI to allow for autophosphorylation.

**Supplementary Figure 3:**
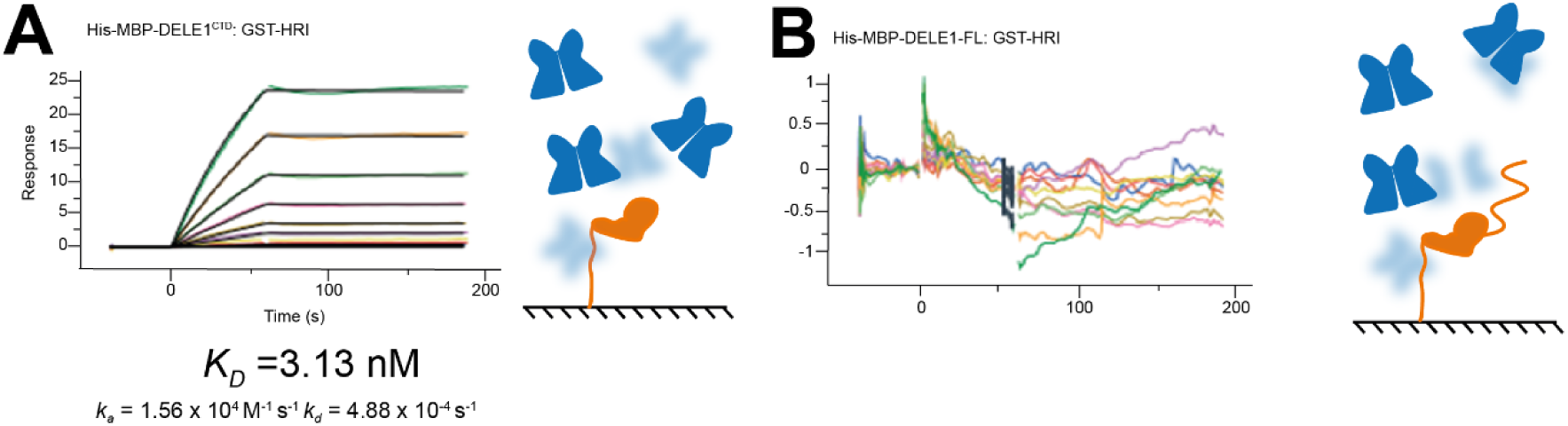
SPR sensorgrams. (**A**) His-MBP-DELE1^CTD^ was immobilised via a His-antibody onto the chip surface. GST-HRI was in the mobile phase in an 8-point concentration curve. An affinity constant could be extracted (**B**) Full-length DELE1 was immobilised in the same manner as (A). No binding of GST-HRI could be detected.

**Supplementary Figure 4:**
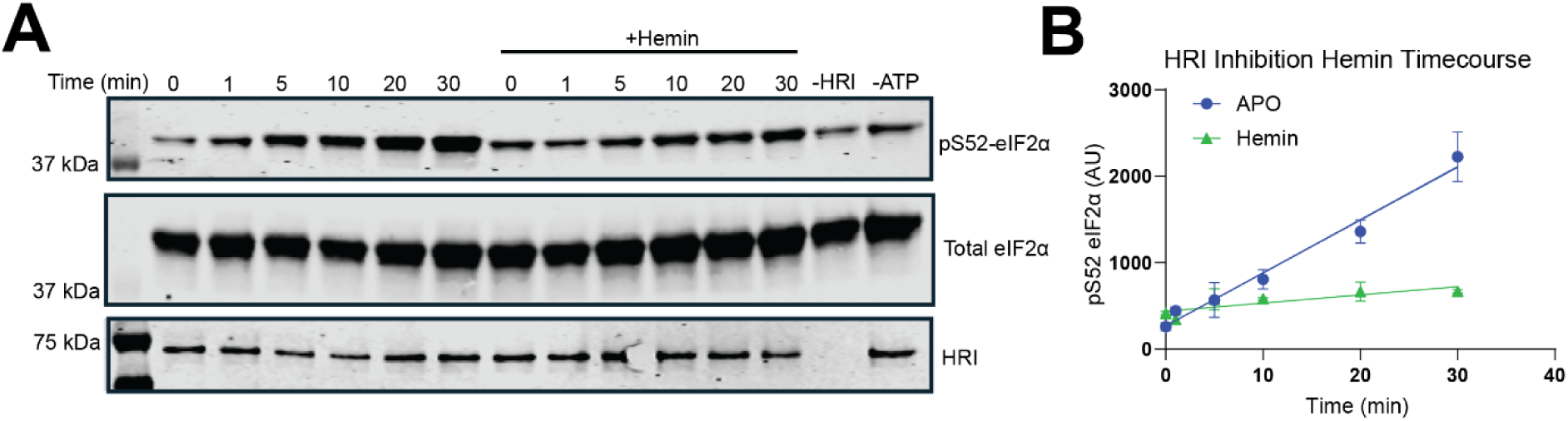
Time course of Hemin inhibition of HRI. (**A**) Western immunoblot analysis of hemin inhibition. 25 nM HRI was incubated with 10 μM ATP for stated times at 25 °C with or without 15 μM hemin. (**B**) Quantification of 3 independent repeats, points are averages with standard deviation as error bars.

**Supplementary Figure 5:**
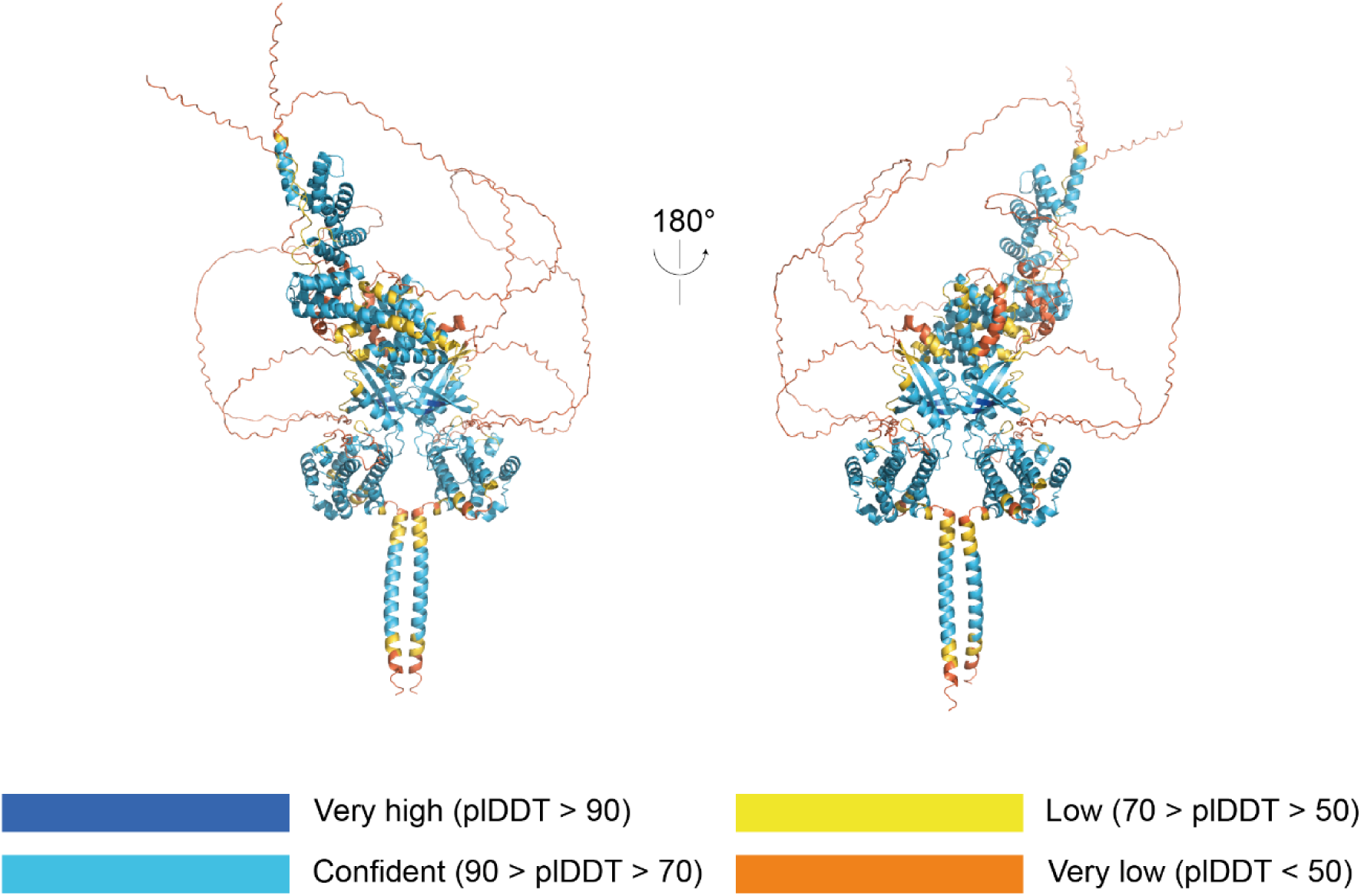
AlphaFold3 prediction of the HRI:DELE1^CTD^ complex. AlphaFold3 model consisting of 2 copies of full-length human HRI (1-630) and 1 copy of DELE1^CTD^ (225-515). ipTM = 0.49, pTM = 0.52 (predicted template modelling (pTM) and interface predicted template modelling (ipTM)).

**Supplementary Table 1.0:** Details of Constructs. All constructs are of human genes.

| Protein | Expressed Protein | Vector Type | Plasmid | Identifier |
| --- | --- | --- | --- | --- |
| eIF2α (EIF2S1) | His-Tev-eIF2α | Bacterial | pOPTH(tev) | pOP819 |
| HRI (EIF2AK1) | His-TEV-HRI | Bacterial | pET15 | DU78201 |
| HRI (EIF2AK1) | GST-TEV-HRI | Bacterial | pGEX6P1 | DU67073 |
| DELE1 (DELE1) | His-MBP-DELE1(225-515) | Bacterial | pMEX3C | DU77970 |
| DELE1 (DELE1) | His-MBP-DELE1 | Bacterial | pMEX3C | DU77964 |

## References

Abdel-Nour M, Carneiro LAM, Downey J, Tsalikis J, Outlioua A, Prescott D, Costa LSD, Hovingh ES, Farahvash A, Gaudet RG, et al (2019) The heme-regulated inhibitor is a cytosolic sensor of protein misfolding that controls innate immune signaling. Science 365

Abramson J, Adler J, Dunger J, Evans R, Green T, Pritzel A, Ronneberger O, Willmore L, Ballard AJ, Bambrick J, et al (2024) Accurate structure prediction of biomolecular interactions with AlphaFold 3. Nature: 1–3

Bi PY, Killackey SA, Schweizer L, Arnoult D, Philpott DJ & Girardin SE (2024) Cytosolic retention of HtrA2 during mitochondrial protein import stress triggers the DELE1-HRI pathway. Commun Biol 7: 391

Bora P, Zaman M, Oviedo S, Kutseikin S, Madrazo N, Mathur P, Pannikkat M, Krasny S, Aldakhlallah R, Chu A, et al (2025) Drug repurposing screen identifies an HRI activating compound that promotes adaptive mitochondrial remodeling in MFN2-deficient cells. Proc Natl Acad Sci 122: e2517552122

Cao M & Masson GR (2026) Structural insights into HRI kinase activity and inhibition. Biochem Soc Trans 54: 561–569

Chen J-J (2025) HRI protein kinase in cytoplasmic heme sensing and mitochondrial stress response: relevance to hematological and mitochondrial diseases. J Biol Chem: 108494

Crosby JS, Lee K, London IM & Chen J-J (1994) Erythroid Expression of the Heme-Regulated eIF-2α Kinase. Mol Cell Biol 14: 3906–3914

Farrell PJ, Balkow K, Hunt T, Jackson RJ & Trachsel H (1977) Phosphorylation of initiation factor eIF-2 and the control of reticulocyte protein synthesis. Cell 11: 187–200

Fessler E, Eckl E-M, Schmitt S, Mancilla IA, Meyer-Bender MF, Hanf M, Philippou-Massier J, Krebs S, Zischka H & Jae LT (2020) A pathway coordinated by DELE1 relays mitochondrial stress to the cytosol. Nature: 1–5

Guo X, Aviles G, Liu Y, Tian R, Unger BA, Lin Y-HT, Wiita AP, Xu K, Correia MA & Kampmann M (2020) Mitochondrial stress is relayed to the cytosol by an OMA1–DELE1–HRI pathway. Nature: 1–6

Igarashi J, Murase M, Iizuka A, Pichierri F, Martinkova M & Shimizu T (2008) Elucidation of the Heme Binding Site of Heme-regulated Eukaryotic Initiation Factor 2α Kinase and the Role of the Regulatory Motif in Heme Sensing by Spectroscopic and Catalytic Studies of Mutant Proteins*. J Biol Chem 283: 18782–18791

Igarashi J, Sasaki T, Kobayashi N, Yoshioka S, Matsushita M & Shimizu T (2011) Autophosphorylation of heme-regulated eukaryotic initiation factor 2α kinase and the role of the modification in catalysis. FEBS J 278: 918–928

Inglis AJ, Masson GR, Shao S, Perisic O, McLaughlin SH, Hegde RS & Williams RL (2019) Activation of GCN2 by the ribosomal P-stalk. Proc National Acad Sci 116: 201813352

Kanta S, Vinciauskaite V, Neill G, Muqit M & Masson G (2025) Structural Insights into allosteric inhibition of HRI kinase by heme binding via HDX-MS. Biochem J

Masson GR, Burke JE, Ahn NG, Anand GS, Borchers C, Brier S, Bou-Assaf GM, Engen JR, Englander SW, Faber J, et al (2019) Recommendations for performing, interpreting and reporting hydrogen deuterium exchange mass spectrometry (HDX-MS) experiments. Nature methods 16: 595–602

Peterle D, Wales TE & Engen JR (2022) Simple and Fast Maximally Deuterated Control (maxD) Preparation for Hydrogen–Deuterium Exchange Mass Spectrometry Experiments. Analytical chemistry 94: 10142–10150

Rafie-Kolpin M, Chefalo PJ, Hussain Z, Hahn J, Uma S, Matts RL & Chen J-J (2000) Two Heme-binding Domains of Heme-regulated Eukaryotic Initiation Factor-2α Kinase N TERMINUS AND KINASE INSERTION*. J Biol Chem 275: 5171–5178

Rafie-Kolpin M, Han A-P & Chen J-J (2003) Autophosphorylation of Threonine 485 in the Activation Loop Is Essential for Attaining eIF2α Kinase Activity of HRI †. Biochemistry 42: 6536–6544

Rajasekaran MB, Booth J, Crepin DF, Roe SM, Zhou L, Hussain R, Gianga T-M, Siligardi G, Gonzalez-Mendez R, Staikopoulou M, et al (2026) Structure-function studies of HRIKD-ΔKI, a Minimal Kinase Domain of Human Heme-Regulated Inhibitor Kinase. bioRxiv: 2026.07.06.735516

Ricketts MD, Emptage RP, Blobel GA & Marmorstein R (2022) The heme-regulated inhibitor kinase requires dimerization for heme-sensing activity. J Biol Chem 298: 102451

Sekine Y, Houston R, Eckl E-M, Fessler E, Narendra DP, Jae LT & Sekine S (2023) A mitochondrial iron-responsive pathway regulated by DELE1. Mol Cell 83: 2059–2076.e6

Singh PK, Agarwal S, Volpi I, Wilhelm LP, Becchi G, Keenlyside A, Macartney T, Toth R, Rousseau A, Masson GR, et al (2025) Kinome screening identifies integrated stress response kinase EIF2AK1/HRI as a negative regulator of PINK1 mitophagy signaling. Sci Adv 11: eadn2528

Vinciauskaite V & Masson GR (2023) Fundamentals of HDX-MS. Essays Biochem 67: 301–314

Yang J, Baron KR, Pride DE, Schneemann A, Guo X, Chen W, Song AS, Aviles G, Kampmann M, Wiseman RL, et al (2023) DELE1 oligomerization promotes integrated stress response activation. Nat Struct Mol Biol 30: 1295–1302

Yang JM, London IM & Chen JJ (1992) Effects of hemin and porphyrin compounds on intersubunit disulfide formation of heme-regulated eIF-2 alpha kinase and the regulation of protein synthesis in reticulocyte lysates. J Biol Chem 267: 20519–20524

Yang M, Mo Z, Walsh K, Liu W, Irahal IN, Arnoult D & Guo X (2026) The integrated stress response suppresses PINK1-dependent mitophagy by preserving mitochondrial import efficiency. Nat Commun

Zebrucka KP, Koryga I, Mnich K, Ljujic M, Samali A & Gorman AM (2016) The integrated stress response. EMBO reports 17: 1374–1395

Zhang X, Schuler M-H, Çetin G, Eckl E-M, Rheinemann L, Mergner J, Steigenberger B, Pichlmair A & Jae LT (2026) An ancient mitochondrial program tunes translation to haem availability. Nature: 1–8

