## Supplementary material for "Structural Basis for Activation of HRI by the DELE1 C-terminal domain": DELE1_HDX

226-242: LSLEEAVTSIQQLFQLS (#1)

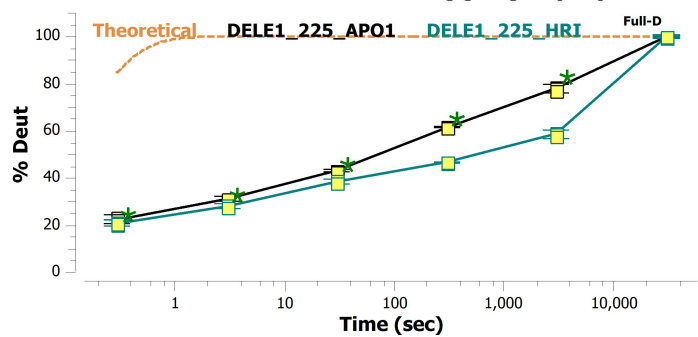

228-240: LEEAVTSIQQLFQ (#2)

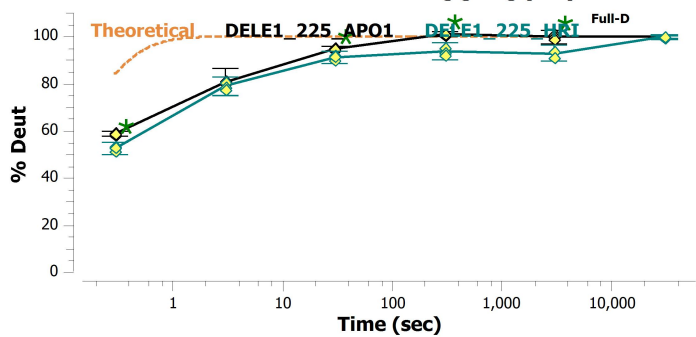

229-241: EEA VTSIQQLFQL (#3)

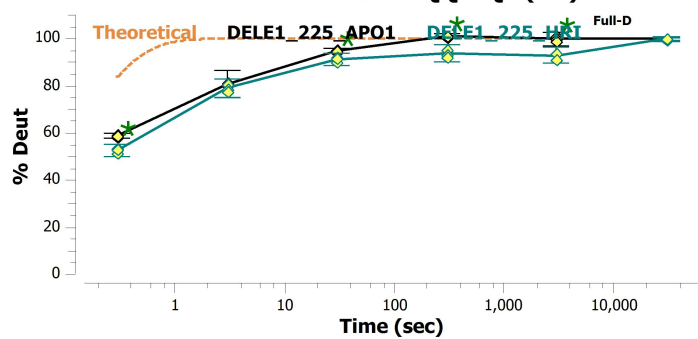

230-241: EAVTSIQQLFQL (#4)

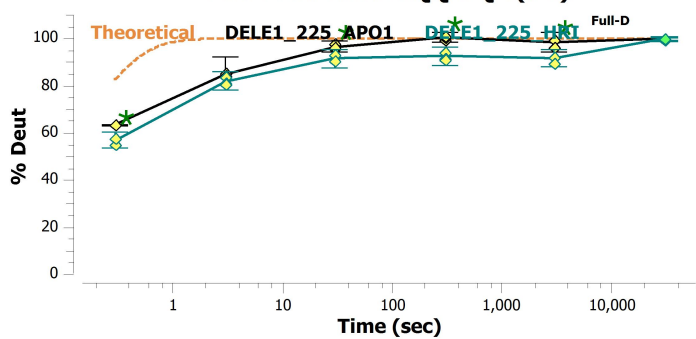

231-241: AVTSIQQLFQL (#5)

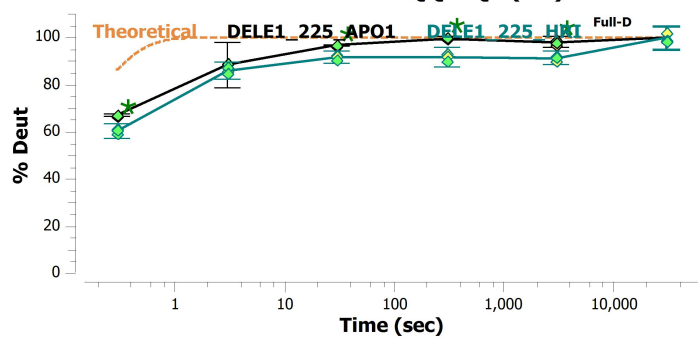

242-247: SVSIAF (#6)

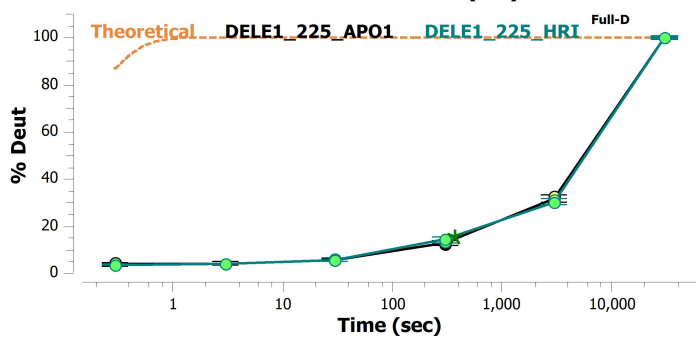

247-255: FNFLGTENM (#7)

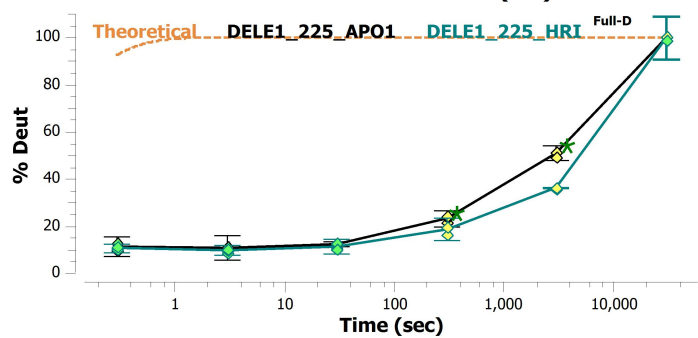

247-263: FNFLGTENMKSGDHTAA (#8)

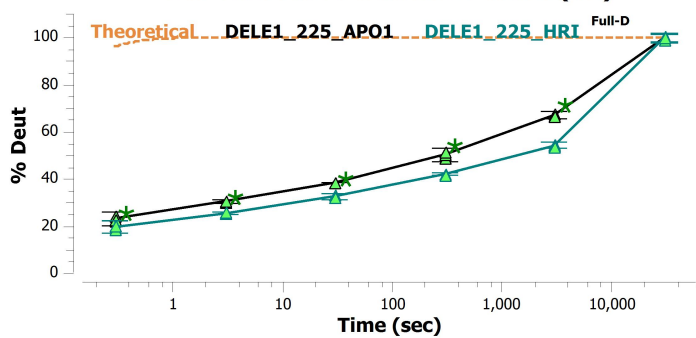

247-264: FNFLGTENMKSGDHTAAF (#9)

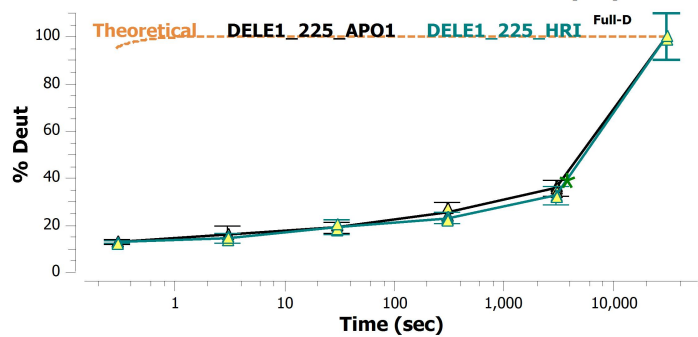

248-255: NFLGTENM (#10)

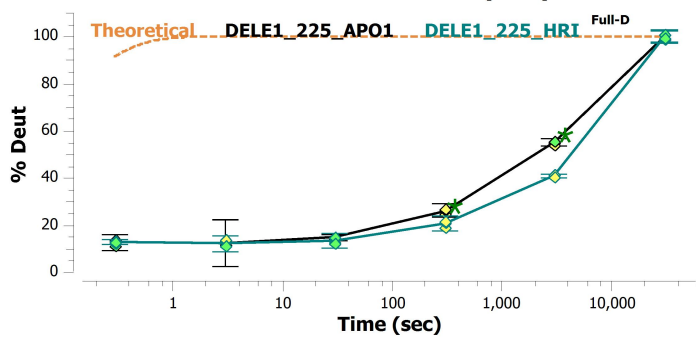

248-257: NFLGTENMK5 (#11)

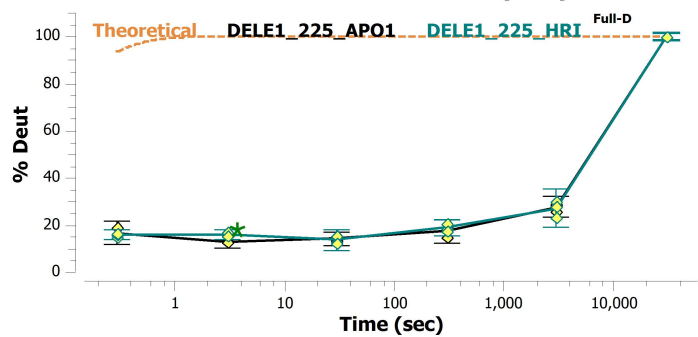

248-263: NFLGTENMK5GDHTAA (#12)

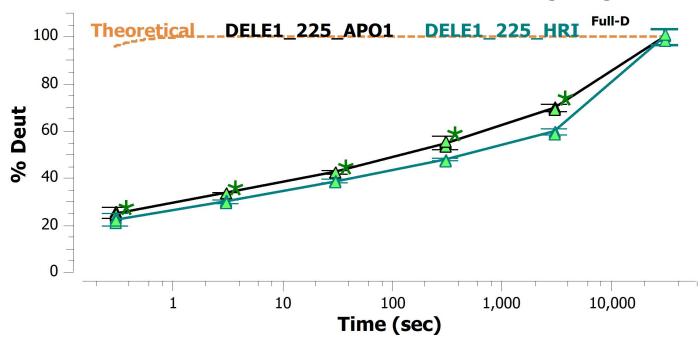

248-264: NFLGTENMK5GDHTAAF (#13)

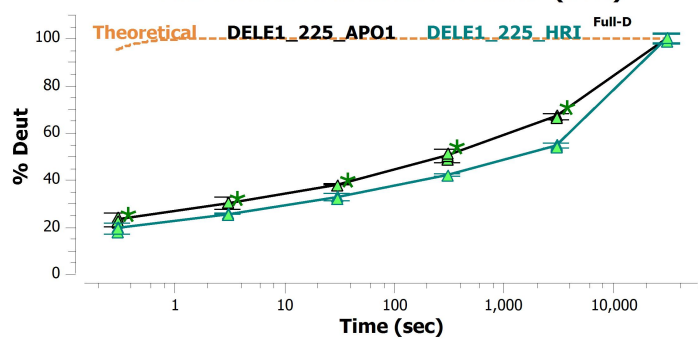

248-265: NFLGTENMK5GDHTAAFS (#14)

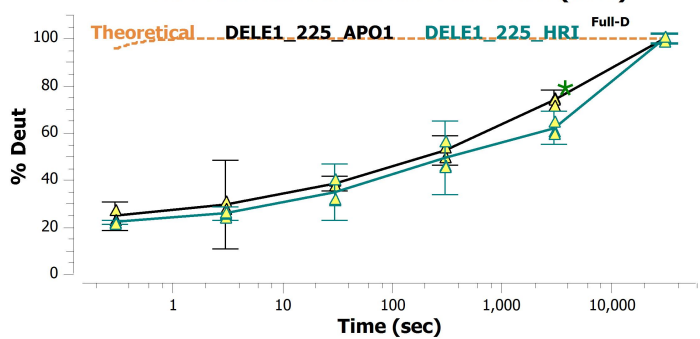

249-263: FLGTENMK5GDHTAA (#15)

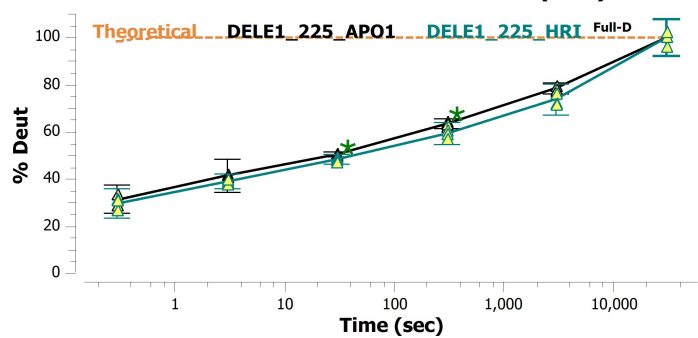

249-264: FLGTENMK5GDHTAAF (#16)

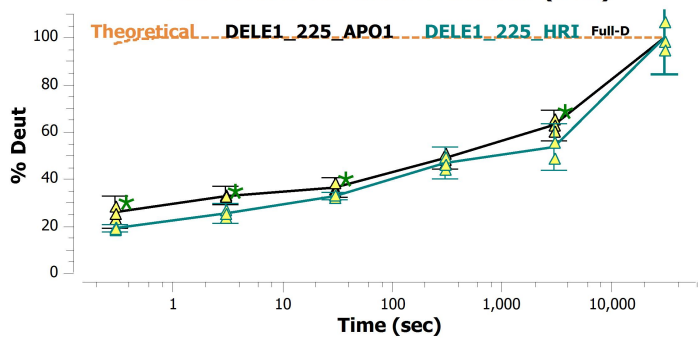

250-261: LGTENMK5GDHT (#17)

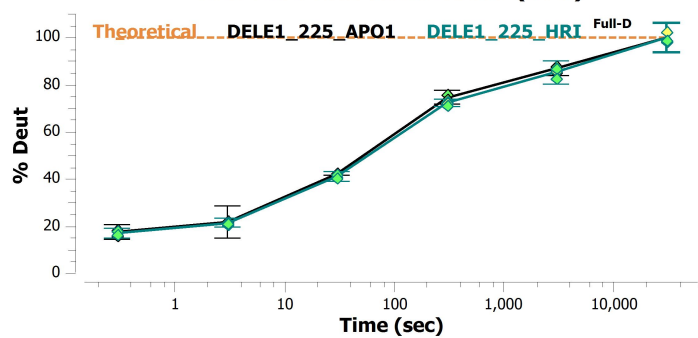

250-263: LGTENMK5GDHTAA (#18)

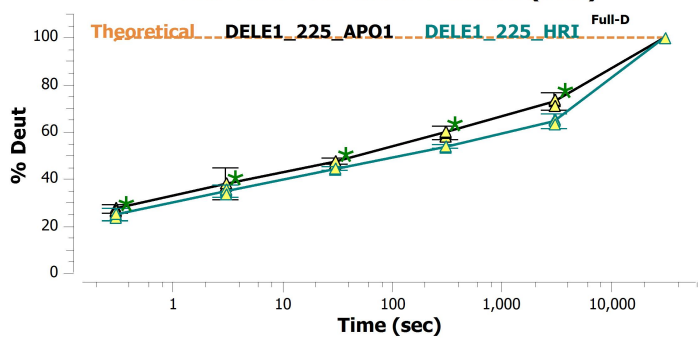

250-264: LGTENMK5GDHTAAF (#19)

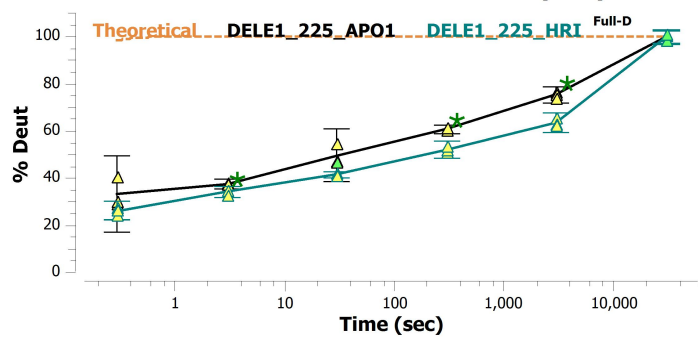

250-268: LGTENMK5GDHTAAFSYFQ (#20)

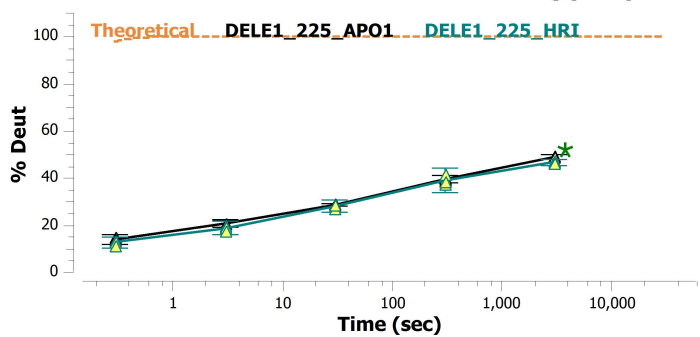

251-263: GTENMKSGDHTAA (#21)

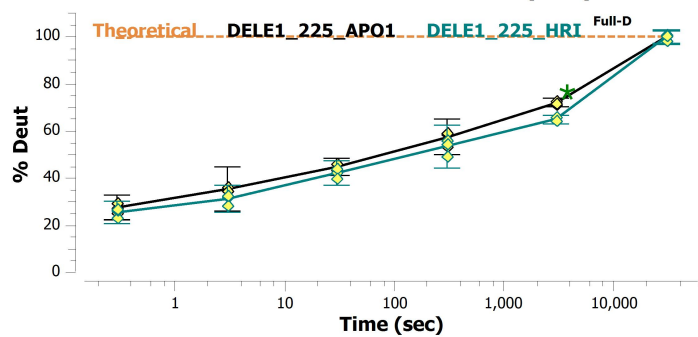

256-264: KSGDHTAAF (#22)

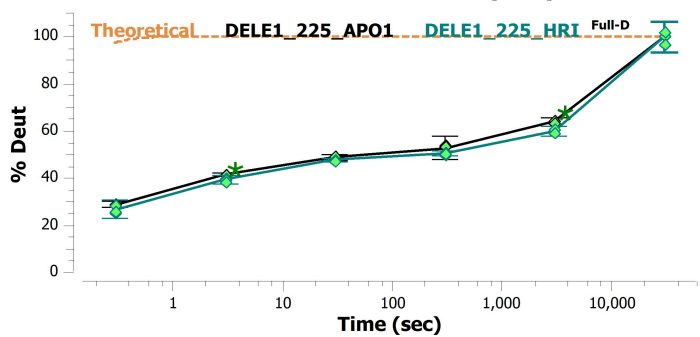

256-265: KSGDHTAAFS (#23)

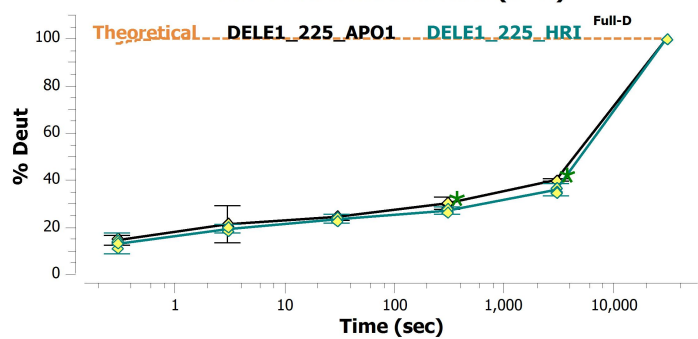

265-274: SYFQKAAARG (#24)

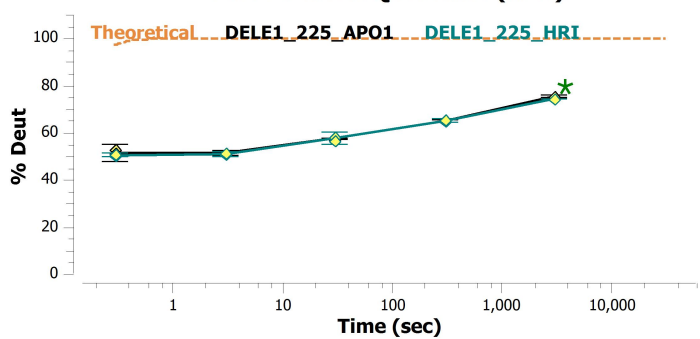

265-301:
SYFQKAAARGYSKAQYNAGLCHEHGRGTPRDISKAVL
(#25)

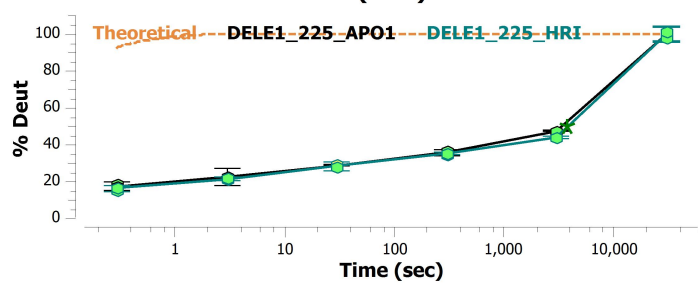

268-281: QKAAARGYSKAQYN (#26)

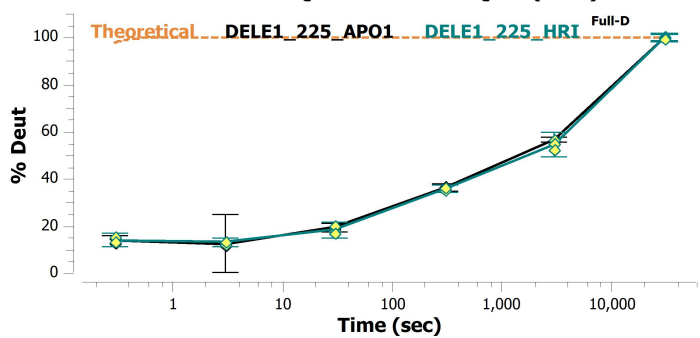

276-314:
SKAQYNAGLCHEHGRGTPRDISKAVLYYQLAASQGHSLA
(#27)

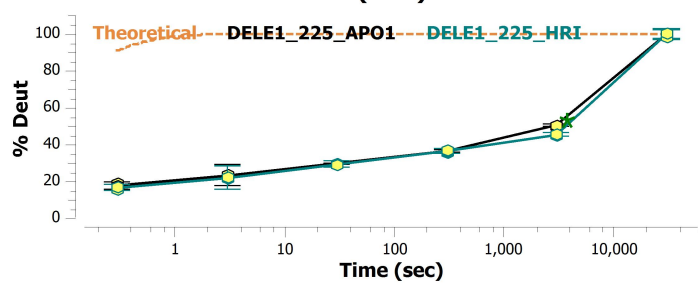

284-301: LCHEHGRGTPRDISKAVL (#28)

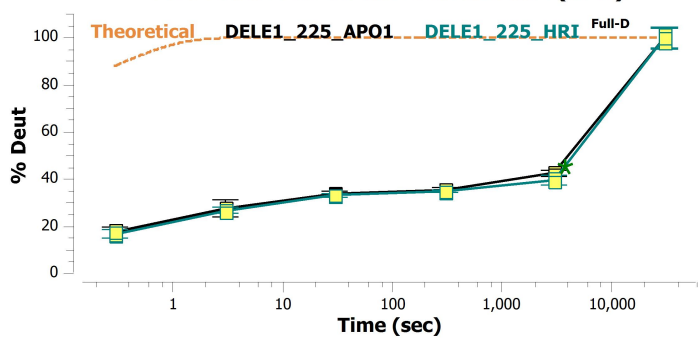

285-301: CHEHGRGTPRDISKAVL (#29)

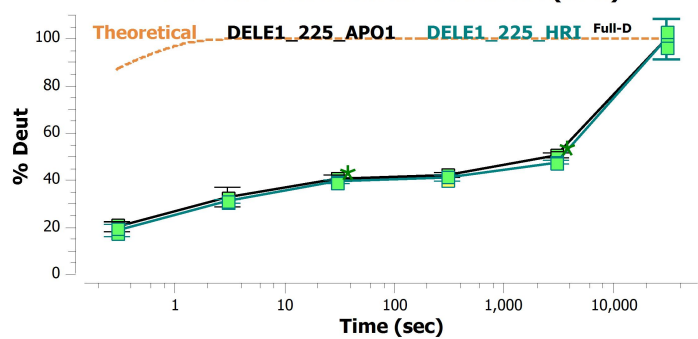

293-305: PRDISKAVLYYQL (#30)

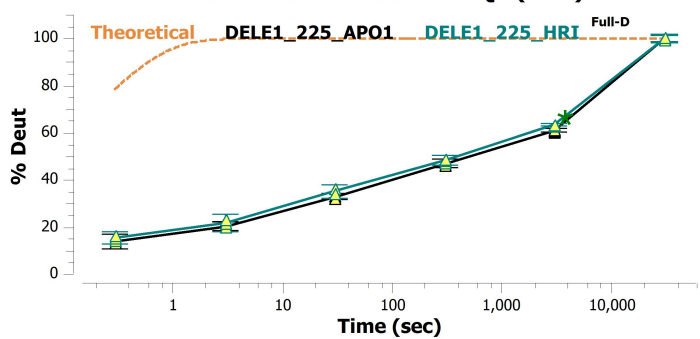

299-312: AVLYYQLAASQGHGS (#31)

300-314: VLYYQLAASQGHSLA (#32)

302-313: YYQLAASQGHSL (#33)

302-315: YYQLAASQGHSLAQ (#34)

302-316: YYQLAASQGHSLAQY (#35)

302-318: YYQLAASQGHSLAQYRY (#36)

302-321: YYQLAASQGHSLAQYRYARC (#37)

302-322: YYQLAASQGHSLAQYRYARCL (#38)

303-315: YQLAASQGHSLAQ (#39)

303-316: YQLAASQGHSLAQY (#40)

303-318: YQLAASQGHSLAQYRY (#41)

303-321: YQLAASQGHSLAQYRYARC (#42)

303-322: YQLAASQGHSLAQYRYARCL (#43)

304-317: QLAASQGHSLAQYR (#44)

306-315: AASQGHSLAQ (#45)

306-316: AASQGHSLAQY (#46)

306-318: AASQGHSLAQYRY (#47)

306-321: AASQGHSLAQYRYARC (#48)

306-322: AASQGHSLAQYRYARCL (#49)

308-321: SQGHSLAQYRYARC (#50)

316-321: YRYARC (#51)

318-337: YARCLLRDPASSWNPERQRA (#52)

322-335: LLRDPASSWNPERQ (#53)

322-337: LLRDPASSWNPERQRA (#54)

322-339: LLRDPASSWNPERQRAVS (#55)

322-340: LLRDPASSWNPERQRAVSL (#56)

322-341: LLRDPASSWNPERQRAVSL (#57)

323-340: LRDPASSWNPERQRAVSL (#58)

323-341: LRDPASSWNPERQRAVSL (#59)

327-356: ASSWNPERQRAVSLKQAADSGLREAQAF (#60)

328-337: SSWNPERQRA (#61)

328-340: SSWNPERQRAVSL (#62)

338-356: VSLLKQAADSGRLREAQAF (#63)

341-348: LKQAADSG (#64)

341-353: LKQAADSGRLREAQ (#65)

341-354: LKQAADSGRLREAQA (#66)

341-355: LKQAADSGRLREAQAF (#67)

341-356: LKQAADSGRLREAQAF (#68)

345-363: ADSGLREAQAFGLVLTKE (#69)

350-359: REAQAFGLVL (#70)

360-377: FTKEPYLDEQRAVKYLWL (#81)

367-374: DEQRAVKY (#82)

367-377: DEQRAVKYLWL (#83)

368-374: EQRAVKY (#84)

369-377: QRAVKYLWL (#85)

370-377: RAVKYLWL (#86)

375-393: LWLAANNQDSQSRVHLGIC (#87)

375-394: LWLAANNQDSQSRVHLGICY (#88)

376-393: WLAANNQDSQSRVHLGIC (#89)

377-393: LAANNQDSQSRVHLGIC (#90)

378-393: AANNGDSQSRVHLGIC (#91)

378-394: AANNGDSQSRVHLGICY (#92)

380-393: NNGDSQSRVHLGIC (#93)

382-395: GDSQSRVHLGICYE (#94)

391-417: GICYEKLGVQRNLGEALRCYQSSAAL (#95)

394-406: YEKLVGVQRNLGE (#96)

394-407: YEKLVGVQRNLGEA (#97)

394-408: YEKLVGVQRNLGEAL (#98)

394-410: YEKLVGVQRNLGEALRC (#99)

395-407: EKLVGVQRNLGEA (#100)

395-410: EKGLGVQRNLGEALRC (#101)

406-423: EALRCYQQSAALGNEAAQ (#102)

409-417: RCYQQAAL (#103)

411-423: YQQAALGNEAAQ (#104)

411-424: YQQAALGNEAAQE (#105)

411-426: YQQAALGNEAAQERL (#106)

411-428: YQQAALGNEAAQERLRA (#107)

411-429: YQQAALGNEAAQERLRAL (#108)

411-430: YQQAALGNEAAQERLRALF (#109)

411-433: YQQAALGNEAAQERLRALFSMG (#110)

425-442: RLRLFSMGAAAPGPSDL (#111)

427-442: RALFSMGAAAPGPSDL (#112)

429-442: LFSMGAAAPGPSDL (#113)

430-442: FSMGAAAPGPSDL (#114)

431-442: SMGAAAPGPSDL (#115)

448-455: KSFSSPSL (#116)

453-475: PSLCSLNTLLAGTSRLPHASSTG (#117)

456-475: CSLNTLLAGTSRLPHASSTG (#118)

458-479: LNTLLAGTSRLPHASSTGNLGL (#119)

459-479: NTLLAGTSRLPHASSTGNLGL (#120)

459-480: NTLLAGTSRLPHASSTGNLGLL (#121)

462-479: LAGTSRLPHASSTGNLGL (#122)

462-480: LAGTSRLPHASSTGNLGLL (#123)

463-479: AGTSRLPHASSTGNLGL (#124)

463-480: AGTSRLPHASSTGNLGLL (#125)

463-492: AGTSRLPHASSTGNLGLLCRSGHLGASLEA (#126)

479-496: LLCRSGHLGASLEASSRA (#127)

480-490: LCRSGHLGASL (#128)

481-490: CRSGHLGASL (#129)

481-501: CRSGHLGASLEASSRAIPHP (#130)

484-497: GHLGASLEASSRAI (#131)

490-510: LEASSRAIPHPYPLERSVVR (#132)

491-505: EASSRAIPHPYPLE (#133)

491-507: EASSRAIPHPYPLERS (#134)

493-514: SSRAIPHPYPLERSVRLGFG (#135)

495-505: RAIPHPYPLE (#136)

495-507: RAIPHPYPLERS (#137)

506-514: RSVRLGFG (#138)
