## Supplementary material for "Structural Basis for Activation of HRI by the DELE1 C-terminal domain": HRI_HDX

**1-29: MQGGNSGVRKREEEGDGAGAVAAPPAIDF (#1)**

**1-46: GNSGVRKREEEGDGAGAVAAPPAIDFPAEGPDPEYDESDI (#2)**

**44-57: PAEIQLKEPLQQP (#3)**

**47-59: IQVLKEPLQQPTF (#4)**

**47-62: IQVLKEPLQQPTFPFA (#5)**

**47-64: IQVLKEPLQQPTFPFAVA (#6)**

**47-66: IQVLKEPLQQPTFPFAVANQ (#7)**

**47-67: IQVLKEPLQQPTFPFAVANQL (#8)**

**47-68: IQVLKEPLQQPTFPFAVANQLL (#9)**

**49-66: VLKEPLQQPTFPFAVANQ (#10)**

73-92: LEHLSHVHEPNPLRSRQVFK (#21)

73-93: LEHLSHVHEPNPLRSRQVFKL (#22)

73-97: LEHLSHVHEPNPLRSRQVFKLLCQT (#23)

74-93: EHLSHVHEPNPLRSRQVFKL (#24)

86-106: RSRQVFKLLCQTFIKMGLLSS (#25)

98-104: FIKMGLL (#26)

98-106: FIKMGLLSS (#27)

98-107: FIKMGLLSSF (#28)

99-106: IKMGLLSS (#29)

99-107: IKMGLLSSF (#30)

**99-127: IKMGLLSSFTCSDEFSSRLHHNRAITHL (#31)**

**100-106: KMGLLSS (#32)**

**108-127: TCSDEFSSRLHHNRAITHL (#33)**

**111-140: DEFSSRLHHNRAITHLMRSAKERVQRDPC (#34)**

**113-127: FSSRLHHNRAITHL (#35)**

**114-127: SSLRLHHNRAITHL (#36)**

**115-127: SRLHHNRAITHL (#37)**

**117-127: RLHHNRAITHL (#38)**

**117-142: RLHHNRAITHLMRSAKERVQRDPCED (#39)**

**127-155: LMRSKERVQRDPCEDISRIQKIRSREVA (#40)**

128-141: MRS AKERVQRDPCE (#41)

128-142: MRS AKERVQRDPCE D (#42)

128-153: MRS AKERVQRDPCE DISRIQKIRSRE (#43)

141-152: EDISRIQKIRS R (#44)

142-156: DISRIQKIRSREVAL (#45)

143-153: ISRIQKIRSRE (#46)

143-156: ISRIQKIRSREVAL (#47)

144-153: SRIQKIRSRE (#48)

144-156: SRIQKIRSREVAL (#49)

154-166: VALEAQT SRYLNE (#50)

156-166: LEAQTSRYLNE (#51)

157-166: EAQTSRYLNE (#52)

157-167: EAQTSRYLNEF (#53)

158-166: AQTSRYLNE (#54)

170-192: LAILGKGGYGRVYKVRNKLDGQY (#55)

171-181: AILGKGGYGRV (#56)

171-192: AILGKGGYGRVYKVRNKLDGQY (#57)

173-192: LGKGGYGRVYKVRNKLDGQY (#58)

193-208: YAIKKILIKGATKTVC (#59)

193-209: YAIKKILIKGATKTVCN (#60)

**194-209: AIKKILIKGATKTVCM (#61)**

**209-229: MKVLREVKVLAGLQHPNIVGY (#62)**

**210-218: KVLREVKVL (#63)**

**210-229: KVLREVKVLAGLQHPNIVGY (#64)**

**210-233: KVLREVKVLAGLQHPNIVGYHTAW (#65)**

**210-251: VLREVKVLAGLQHPNIVGYHTAWIEHVVHIQPRADRAAI (#66)**

**211-246: VLREVKVLAGLQHPNIVGYHTAWIEHVVHIQPRADR (#67)**

**215-229: VKVLAGLQHPNIVGY (#68)**

**215-233: VKVLAGLQHPNIVGYHTAW (#69)**

**215-251: VKVLAGLQHPNIVGYHTAWIEHVVHIQPRADRAAIEL (#70)**

**281-310: FAEPTPEKEKRFGESDTENQNNKSVKYTTN (#81)**

**281-311: FAEPTPEKEKRFGESDTENQNNKSVKYTTNL (#82)**

**282-295: AEPTPEKEKRFGES (#83)**

**282-310: AEPTPEKEKRFGESDTENQNNKSVKYTTN (#84)**

**282-311: AEPTPEKEKRFGESDTENQNNKSVKYTTNL (#85)**

**296-311: DTENQNNKSVKYTTNL (#86)**

**303-318: KSVKYTTNLVIRESGE (#87)**

**304-340: SVKYTTNLVIRESGELESTLEQLQENGLAGLSASSIVE (#88)**

**311-319: LVIRESGEL (#89)**

**311-324: LVIRESGELESTLE (#90)**

**325-351: LQENGLAGLSASSIVEQQLPLRRNSHL (#101)**

**325-355: LQENGLAGLSASSIVEQQLPLRRNSHLEESF (#102)**

**331-351: AGLSASSIVEQQLPLRRNSHL (#103)**

**334-351: SASSIVEQQLPLRRNSHL (#104)**

**334-354: SASSIVEQQLPLRRNSHLEES (#105)**

**334-355: SASSIVEQQLPLRRNSHLEESF (#106)**

**338-351: IVEQQLPLRRNSHL (#107)**

**338-353: IVEQQLPLRRNSHLEE (#108)**

**338-354: IVEQQLPLRRNSHLEES (#109)**

**341-351: QQLPLRRNSHL (#110)**

395-411: IVERNKRGREYVDESAC (#121)

405-415: YVDESACPYVM (#122)

416-425: ANVATKIFQE (#123)

416-426: ANVATKIFQEL (#124)

416-428: ANVATKIFQELVE (#125)

417-425: NVATKIFQE (#126)

417-426: NVATKIFQEL (#127)

417-428: NVATKIFQELVE (#128)

418-426: VATKIFQEL (#129)

419-426: ATKIFQEL (#130)

**427-446: VEGVFIHNMGIVHRDLKPR (#131)**

**431-461: FYIHNMGIVHRDLKPRNIFLHGPDQQVKIGD (#132)**

**432-447: YIHNMGIVHRDLKPRN (#133)**

**432-450: YIHNMGIVHRDLKPRNIFL (#134)**

**432-456: YIHNMGIVHRDLKPRNIFLHGPDQQ (#135)**

**432-464: YIHNMGIVHRDLKPRNIFLHGPDQQVKIGDFGL (#136)**

**432-466: YIHNMGIVHRDLKPRNIFLHGPDQQVKIGDFGLAC (#137)**

**433-447: IHNMGIVHRDLKPRN (#138)**

**440-447: HRDLKPRN (#139)**

**448-464: IFLHGPDQQVKIGDFGL (#140)**

496-513: YASPEQLEGSEYDAKSDM (#161)

503-513: EGSEYDAKSDM (#162)

507-513: YDAKSDM (#163)

521-530: LELFQPFGTE (#164)

521-531: LELFQPFGTEM (#165)

521-532: LELFQPFGTEME (#166)

522-532: ELFQPFGTEME (#167)

524-532: FQPFGTEME (#168)

525-531: QPFGTEM (#169)

526-532: PFGTEME (#170)

**538-559: TGLRTGQLPESLRKRCVPVQAKY (#181)**

**538-574: TGLRTGQLPESLRKRCVPVQAKYIQHLTRRNSSQRPSA (#182)**

**538-577: TGLRTGQLPESLRKRCVPVQAKYIQHLTRRNSSQRPSAIQL (#183)**

**541-560: RTGQLPESLRKRCVPVQAKYI (#184)**

**550-574: RKRCVPVQAKYIQHLTRRNSSQRPSA (#185)**

**550-577: RKRCVPVQAKYIQHLTRRNSSQRPSAIQL (#186)**

**560-574: IQHLTRRNSSQRPSA (#187)**

**560-577: IQHLTRRNSSQRPSAIQL (#188)**

**575-582: IQLLQSEL (#189)**

**583-591: FQNSGNVNL (#190)**

583-593: FQNSGNVNLTL (#191)

592-606: TLQMKIIQEKEIAE (#192)

594-603: QMKIIQEKE (#193)

594-607: QMKIIQEKEIAEL (#194)

594-613: QMKIIQEKEIAELKKQLNL (#195)

596-603: KIIQEKE (#196)

596-605: KIIQEKEIA (#197)

596-606: KIIQEKEIAE (#198)

596-607: KIIQEKEIAEL (#199)

596-613: KIIQEKEIAELKKQLNL (#200)

604-613: IAEKKQLNL (#201)

605-613: AELKKQLNL (#202)

608-613: KKQLNL (#203)

608-614: KKQLNLL (#204)
